# Gene network topologies for heterogeneity restoration during self-renewal

**DOI:** 10.64898/2026.11.16.767011

**Authors:** Suchana Chakravarty, Rija Tonny Christian Ramarolahy, Tian Hong

## Abstract

The ability for a subset of cells to establish a heterogeneous cell population with multi-lineage competency is crucial for the self-renewal capacity of tissues and organs. Restoration of heterogeneous gene expression patterns has been observed in pluripotent and multi-potent stem cells, and the heterogeneity restoration can influence the potential of differentiation into multiple cell types. Similarly, cancer cells can acquire the capacity to establish non-genetic heterogeneity that plays a role in cancer progression and resistance to therapeutics. Specific models of gene regulatory networks have been proposed to explain heterogeneity restoration. However, due to the lack of systematic analysis, it is unclear which network topologies are more efficient for this multicellular function, and what intrinsic and extrinsic factors can determine the optimal choice of network topology. In this work, we use mathematical models describing gene expression of single cells and their underlying population dynamics to examine the influence of network topology on the recovery of multimodal expression patterns originating from a homogeneous population. We found that feedback loops are a key factor contributing to heterogeneity restoration. Negative and positive feedback loops support the existence of heterogeneity restoration, but only positive feedback can accelerate the process with low cost of single-cell-level variations of expression. Furthermore, networks with mixed feedback were the most robust ones for heterogeneity restoration with respect to changes in both kinetic rate constants and noise levels. Using experimentally validated network databases, we showed that genes involved in feedback loops are enriched with functions related to cell fate decisions, and we identified examples of realistic gene regulatory networks cell fate dynamics. Our results suggest multiple potential evolutionary paths towards gene circuits governing heterogeneity restoration and key factors contributing to the dynamics of this process. We expect that this systematic study will enable future discoveries of new gene regulatory networks governing progenitor and cancer cell dynamics as well as therapeutic strategies targeting heterogeneity-enabling circuits.

## Introduction

The ability for progenitor cells to self-renew and differentiate into diverse cell types is crucial for development and regeneration. The multi-lineage potential for each of these cells depends on its capacity to efficiently switch its transcriptional program to more than one differentiated cell state with distinct gene expression patterns. It has been shown that clonal cell populations, including embryonic stem cells and multi-potent progenitor cells in mammals, have heterogeneous gene expression patterns even before lineage commitment, and that these patterns play a role in their multi-lineage differentiation potentials ^1–3^. Interestingly, these patterns and their underlying differentiation potentials can be restored from isolated subpopulations with extreme gene expression profiles ^1,4,5^, which can be viewed as a small-scale regeneration process. Similar heterogeneity restoration phenomena were observed in cancer cells that utilize nongenetic heterogeneity to enhance their flexibility of adaptation to multiple microenvironmental conditions ^6^ (**Figure 1**). The behavior, akin to the progenitor cell counterpart, also contributes to resistance to therapeutics ^7,8^, but the gene-network-level mechanism underlying this dynamical property is poorly understood.

**Figure 1.**
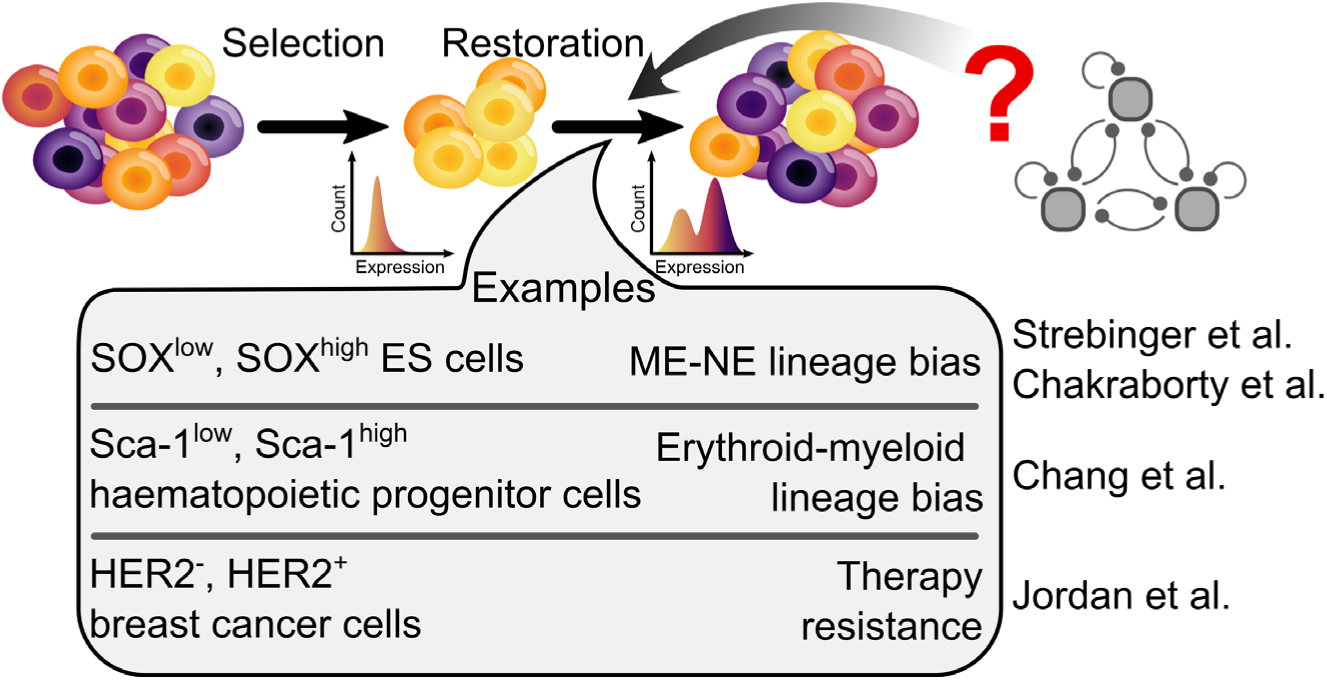
An illustration of heterogeneity restoration in cell populations. A multimodal gene expression pattern is recovered on the timescale of days after selecting a subpopulation with extreme/biased expression from a clonal cell population. This pattern is often associated with diverse lineage potentials or sensitivities to therapeutics.

Two different types of gene regulatory network models have been proposed to explain specific cases of heterogeneity restoration. The first type stems from the observation that heterogeneous gene expression patterns in progenitor and cancer cells are often multimodal. A multimodal distribution under the same condition can be governed by a multistable dynamical system ^4^, which is in turn described by gene regulatory network models containing positive feedback loops, including a widely observed cross-repression motif involving two or more transcription factors ^9,10^. The second type of model is based on robust periodic dynamics of gene expression that can enable the regeneration of diverse gene expression patterns over time. These models often include negative feedback loops that govern oscillations of gene expression without the need of multistability ^11–14^. While both model types have the potential to explain heterogeneity restorations, their stark difference in topology of gene regulatory networks raised the question of whether one type of model has any performance or evolutionary advantage over the other. Furthermore, it is unclear whether other types of models can provide additional dynamical functions to support heterogeneity restoration. Previous computational studies characterized multistate expression patterns produced by a large number of gene networks ^15,16^, but they focused on heterogeneity arising from perturbations of kinetic rate constants rather than dynamics of monoclonal cells, leaving the aforementioned questions concerning collective behaviors of cell populations unaddressed.

In this work, we screened 3325 networks involving one, two or three genes. We used stochastic ordinary differential equations to describe single-cell and population dynamics of progenitor cells starting from extreme gene expression and to examine their ability to restore population-level heterogeneity of single-cell gene expression. We used a series of performance metrics reflecting biological functions, including the existence of heterogeneity restoration, robustness, the speed and the ‘cost’ in terms of overall changes of individual cells’ gene expression. We showed that feedback loops, but not feed-forward loops, significantly contribute to heterogeneity restoration. Negative and positive feedback loops support the existence of heterogeneity restoration with similar strengths, but their effects on the speed are the opposite. We found that mixed feedback networks are more robust than pure feedback networks in heterogeneity restoration with variations of both kinetic rate constants and noise levels. We used statistical analysis with databases of known gene regulations to show that genes involved in feedback loops are enriched for cellular development, cell differentiation and stem cell functions. We identified examples of gene regulatory networks that support experimentally observed heterogeneity recovery in several types of progenitor cells.

Our work relates features of progenitor cell dynamics involving several aspects of heterogeneity restoration to both network topology and concrete gene circuits. Our results suggest multiple paths leading to gene circuits governing heterogeneity restoration. We expect that our systematic analysis will enable future construction of new gene regulatory networks that may be used as regenerative therapeutics, as well as strategies targeting heterogeneity-enabling circuits in cancer cells.

## Results

### Feedback loops promote heterogeneity restoration

We developed a computational framework for simulating cell populations with gene circuits containing up to three nodes (**Figure 2A**). The dynamics of each cell’s gene circuit were described by a system of stochastic ordinary differential equations (SODEs) (see Methods). For each *in silico* experiment, we simulated 500 cells with a uniform initial condition, reflecting an extreme case of an onset of a heterogeneity recovery (HR) right after a selection (e.g. cell sorting) process. We first obtained the time-course distributions of the cell population in terms of the genes’ expression (activity) levels, and we assessed the modality of the distribution at the stationary phase of the simulation (see Methods). Next, the ability for heterogeneity restoration was determined by whether the system can achieve a multimodal distribution of gene expression with well-separated subpopulations. We sampled 3,325 topologically distinct 3-node gene networks. For each network, we randomly sampled 1,000 parameter sets (see Methods. **Supplementary Table S1**). If one or more parameter sets of a network produced multimodal gene expression at stationary phase with two initial conditions at opposite locations of expression space (see Methods. **Figure S1**), then the network was selected as a heterogeneity recovery-enabling (HR-enabling) network (**Figure 2B**).

**Figure 2.**
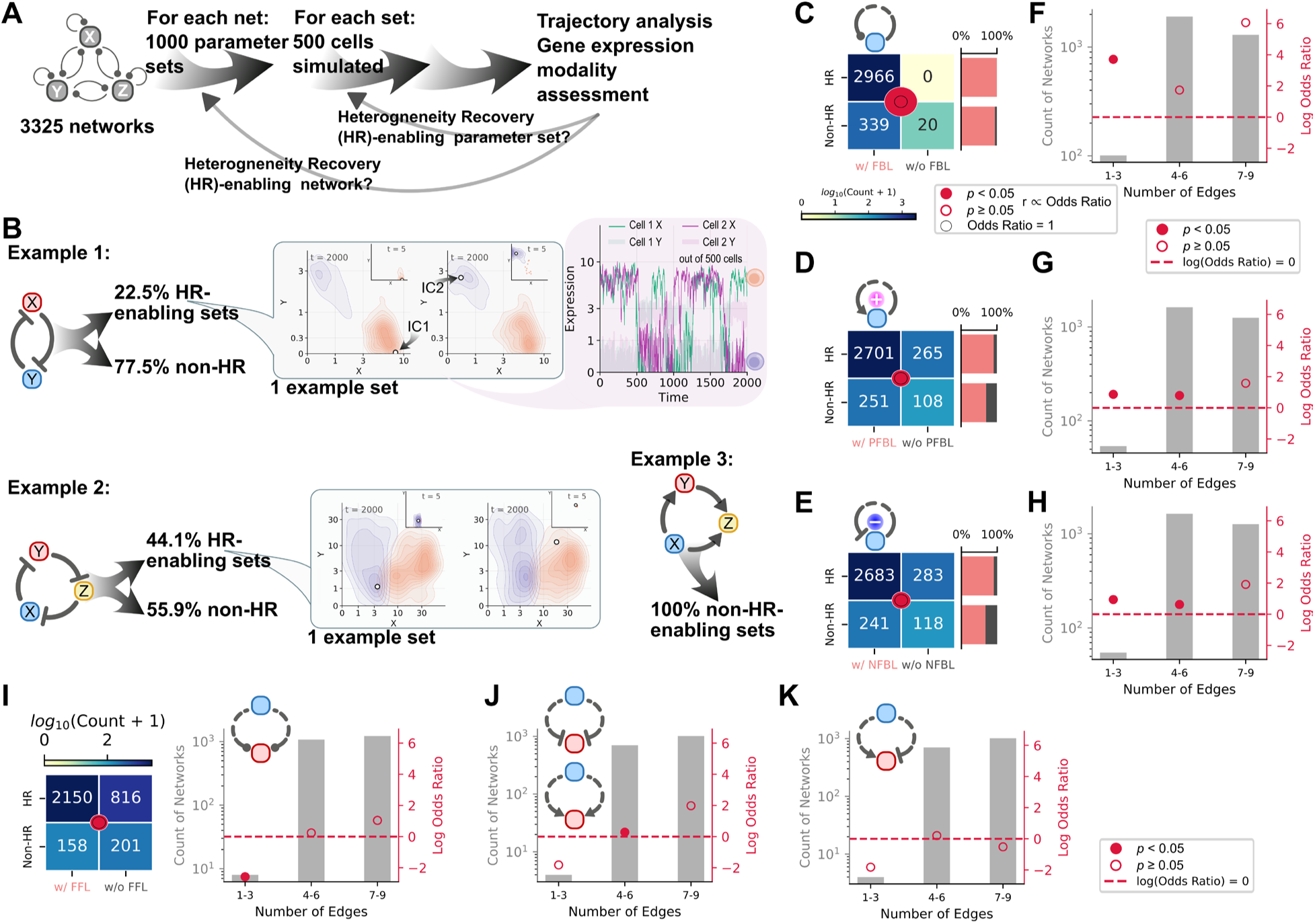
Network topologies that support the existence of heterogeneity recovery. **A**. Illustration of the selection process for HR-enabling parameter sets and networks. **B**. Two examples of HR-enabling networks and one example of non-HR-enabling networks. Gray callouts show distributions of cells at the final and an early (inset) timepoints. Pink callout shows gene expression dynamics in two cells from the population. Expression is shown in the scale of log(x+1). Insets and main plots have the same axis limits. **C-E**. Contingency table for existence of HR and feedback loops (C), positive feedback loops (D) and negative feedback loops (E). **F-H**. Log odds ratios for HR and feedback loops (F), positive feedback loops (G) and negative feedback loops (H) with 3-category stratification of network density. **I-K**. Contingency table for existence of HR and feed-forward loops, and log odds ratios for HR and feed-forward loops (I), coherent feed-forward loops (J), and incoherent feed-forward loops (K). Fisher’s exact test was used for C-K.

Among 3,325 networks, 2966 of them were HR-enabling, and each of these networks contains at least one feedback loop. All of the 20 networks without any feedback loops failed to produce multimodality with any tested parameter set. We observed both the requirement of feedback loop (FBL) for HR and a strong statistical association between feedback loop and HR (**Figure 2C**. Odds ratio (OR) > 4; *p* < 10^−5^, Fisher’s exact test) with the sampled parameter range and the basal gene expression noise. We found that positive feedback loop (PFBL) and negative feedback loop (NFBL) both had positive associations with HR (**Figure 2D** and **E**). Together with those HR-enabling networks that cannot generate bistability (**Figure 2B**, Example 2), the association between NFBL and HR demonstrates that HR is not simply a result of multiple coexisting attractors. In addition, the direction of the associations was consistent across networks with various numbers of edges (network densities), even though the statistical significance was decreased in some categories (**Figure 2F-H**), suggesting that the contributions of feedback loops to HR were not merely due to increasing network density. In contrast to feedback loops, while HR appeared to have a positive overall association with feed-forward loop (FFL), the association was inconsistent and/or insignificant in most density categories of networks, suggesting a ‘Simpson’s paradox’ in which the effect of FFLs was confounded by network density (**Figure 2I-K**). Furthermore, we did not observe the requirement of FFL for achieving HR (**Figure 2I**).

We next used the Mantel–Haenszel (MH) stratification as a more systematic approach to control for the effect of network density measured by the number of edges here (note that the test can only be performed when a stratum contains both HR and non-HR cases) (**Table 1**). As expected, feedback loops contributed to HR positively after MH stratification, whereas the effects of feed-forward loops became negative with the stratification. Overall, positive and negative feedback loops had similar associations with HR, although positive feedback showed weaker statistical significance. Based on the number and the sign of feedback loops, we categorized all networks into 5 types: pure positive (n=381), mixed positive-dominant (n=969), balanced (n=643), mixed negative-dominant (n=959) and pure negative (n=353). We examined HR existence with this finer categorization of feedback compositions, and we found that there was not a significant difference among different proportions of positive/negative feedback loops in their association with HR (**Figure 3A**). These results suggest that the ability to recover multimodal gene expression is not the same as the ability to generate multistability, a unique feature of positive feedback. In addition, we found that HR existence is more likely to be governed by feedback loops involving more than one gene, whereas auto-regulations had overall negative associations with HR for both feedback types (**Figure 3B**). In summary, we found that both positive and negative feedback loops can promote heterogeneity restoration in cell populations, and that feedback loops are required for heterogeneity restoration in general.

**Figure 3.**
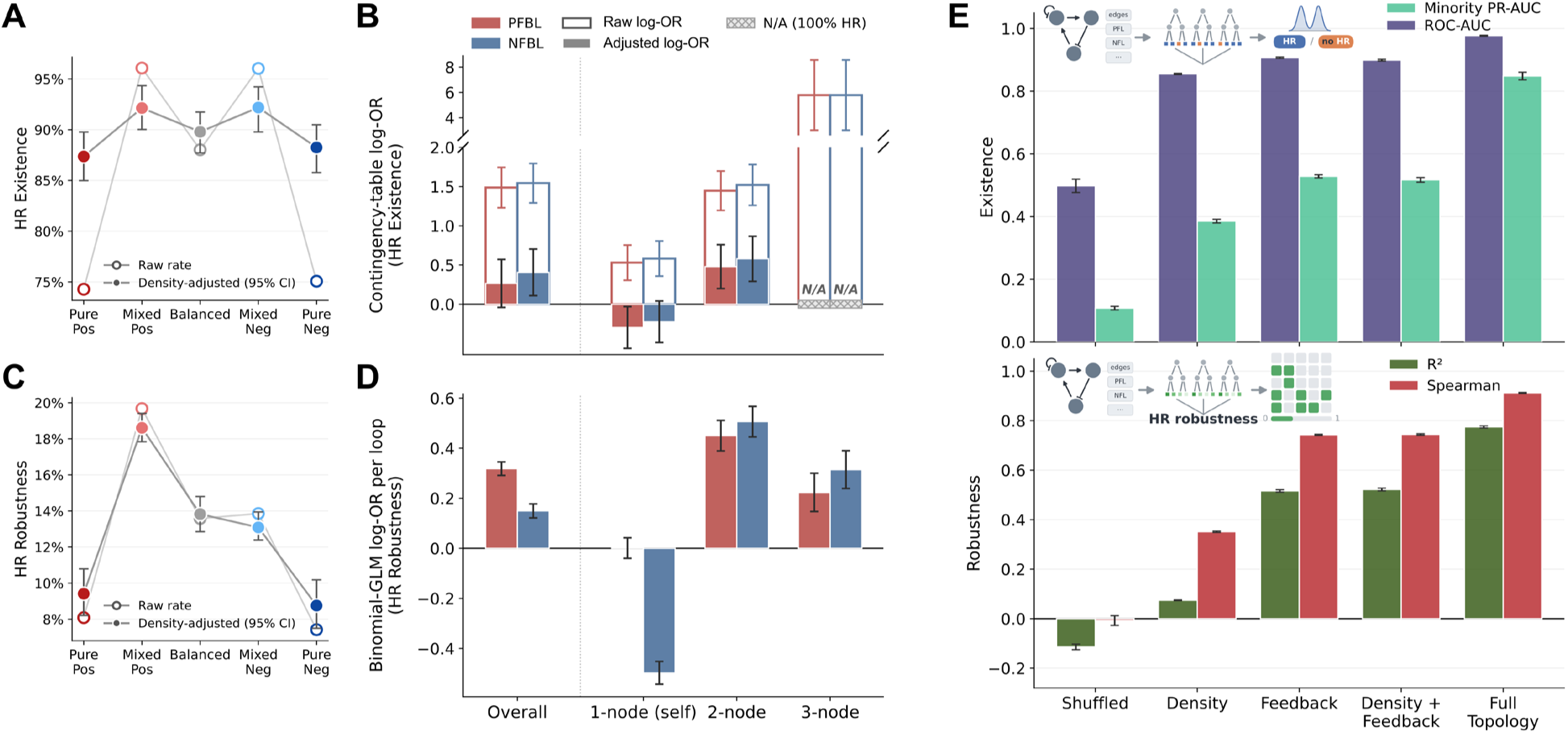
Role of feedback length in HR and predictability of HR with topology. **A**. Fraction of HR-enabling networks in each feedback composition category. Filled circles indicate density-adjusted rate by g-computation from a logistic model in *n*_edge_+ *n*_edge_^2^, with 95% network-bootstrap CIs. **B**. Contingency table log-OR for overall feedback variables (left) and individual feedback length variables (right). Error bars indicate Robins–Breslow–Greenland variance. **C**. Similar to A, but the outcome is the per-network multimodal rate (HR-enabling sets / 1,000), modeled by binomial GLM with network-level bootstrap CIs. **D**. Binomial-GLM log-OR for overall feedback variables and individual feedback length variables model-based. Error bars indicate binomial standard error. **E**. Performance of random forest classifiers (top) and regressors (bottom) with indicated feature groups. ROC-AUC: Receiver Operating Characteristic - Area Under the Curve. PR-AUC: Area Under the Precision-Recall Curve. Minority class refers to non-HR-enabling. 5-fold cross validations were used for each model. Error bars indicate standard errors of the mean.

**Table 1.** Effects of motifs on HR with Mantel–Haenszel (MH) stratification.

| **Motif** | ***n*<sub>network</sub>** | **Raw log-OR** | **MH log-OR [95% CI]** | **Strata** |
|---|---|---|---|---|
| PFBL | 2952 | +1.488 [+1.230, +1.745] | +0.265 [−0.041, +0.571] | 7 (*n*<sub>edge</sub>=1–7) \* |
| NFBL | 2924 | +1.544 [+1.292, +1.795] | +0.406 [+0.109, +0.703] | 7 (*n*<sub>edge</sub>=1–7) \* |
| CFFL | 1722 | +1.221 [+0.975, +1.466] | −0.335 [−0.631, −0.039] | 5 (*n*<sub>edge</sub>=3–7) \*\* |
| IFFL | 1722 | +1.128 [+0.887, +1.369] | −0.481 [−0.779, −0.183] | 5 (*n*<sub>edge</sub>=3–7) \*\* |

\* Edge numbers outside these strata produced either 0% or 100% HR-enabling.
\*\* Edge numbers outside these strata produced either no FFL or 100% HR-enabling.

### Numbers and lengths of feedback loops regulate heterogeneity restoration

Since not all parameter sets (kinetic rate constants) of a network can generate heterogeneity restoration (HR) (**Figure 2B**), we asked whether the two types of feedback loops can influence the percentage of parameter sets that gave rise to HR. This percentage reflects the robustness of a network in generating a multimodal gene expression pattern and its underlying HR. Interestingly, unlike the existence of HR, the compositions of feedback loops had a strong influence on the robustness of HR (**Figure 3C**): mixing the two types of feedback loops increased robustness significantly. This synergy effect seems to be the strongest when the number of positive feedback loops dominates in a network (**Figure 3C**). This result suggests that the two feedback types have distinct mechanisms to support HR, and two types of parameter sets for the same HR-enabling network can complement each other to enhance robustness.

We next used a binomial generalized linear model (GLM) to quantify the influence of the number of feedback loops on the HR percentage/robustness (i.e. the percentage of parameter sets that successfully produced HR). We found that the two types of feedback loops, PFBL and NFBL, both had positive overall influences on robustness when used as model predictors, while the influence from PFBL was stronger than that from NFBL (**Figure 3D**). With a binomial GLM containing 1-node, 2-node, and 3-node feedback loops as predictor variables for HR robustness, we found that the number of negative auto-regulations had a negative effect on HR robustness, whereas the effect of positive auto-regulations was not significant. In contrast, numbers of 2-node and 3-node feedback loops had a definitively positive role in HR robustness (**Figure 3D**). Overall, our results suggest that feedback loops involving more than one gene support HR existence and robustness positively and that mixing feedback types increases HR robustness.

We next asked to what degree one can use network topology features to predict whether a network is HR-enabling. We generated a list of 63 topological features (see **Supplementary Table S2**) and built a random forest classifier model for predicting HR existence. We found that the model achieved excellent performance even for predicting the minority class (non-HR networks) (**Figure 3E**, top right). Since some of these topological features are either trivial (e.g. density) or uninterpretable in terms of biological meaning (e.g. determinant of adjacency matrix), we used a model containing only 6 feedback features representing the occurrences of feedback loops with various lengths (i.e. **Figure 3B**, right). We found that these feedback-related features can provide more than 90% of overall predictability with respect to the full-topology model and more than 60% of predictability for the minority class, a performance significantly better than random and a model with network density feature alone (**Figure 3E**, top). Similarly, a random forest regressor with feedback-related features provided a reasonable predictability for the robustness of HR among HR-enabling networks (**Figure 3E**, bottom). These results suggest that feedback loops and their lengths provide a strong predictive power for the existence and robustness of HR.

### Positive and negative feedback showed distinct heterogeneity restoration dynamics

Our analyses so far have shown a substantial similarity between positive and negative feedback in supporting multimodal gene expression and heterogeneity restoration (HR). While this similarity can be potentially explained by dynamical systems theory in that both types of feedback loops can create nonlinear changes that tend to destabilize a single point attractor, one would still expect that the resulting HR dynamics from the two types of regulatory networks may differ significantly (see more detailed discussion later). We first examined the settling time, defined as the time required for the cell population to remain in a 5% error band in terms of cell fraction (**Figure 4A**). We found that networks with positive feedback had significantly shorter median settling times whereas negative feedback-containing networks had the opposite trend (**Figure 4A**). While this difference is significant, settling times of most HR-enabling parameter sets fell in the range of 1200-1800 time-units (**Figure S2**), corresponding to approximately 7-12 days, a reasonable timeframe for mammalian cells’ HR. To dissect the roles of feedback loops with various lengths in settling time, we built an ordinary least squares (OLS) model with occurrences of 6 types of feedback loops as predictor variables. We found that NFBLs with all 3 lengths slowed down HR, whereas each of the 2-node positive feedback accelerated it (**Figure 4B**). We next asked whether there is a correlation between the robustness of a network’s ability to achieve HR and its median settling time across parameter sets, and we found that the settling time-robustness correlation was positive for mixed but not pure feedback networks (**Figure 4C**, **Figure S3**), suggesting a speed-robustness tradeoff between the two traits in mixed feedback networks. Interestingly, pure positive feedback networks had a negative settling time-robustness correlation, suggesting that HR can be either achieved quickly, or never achieved in this category.

**Figure 4.**
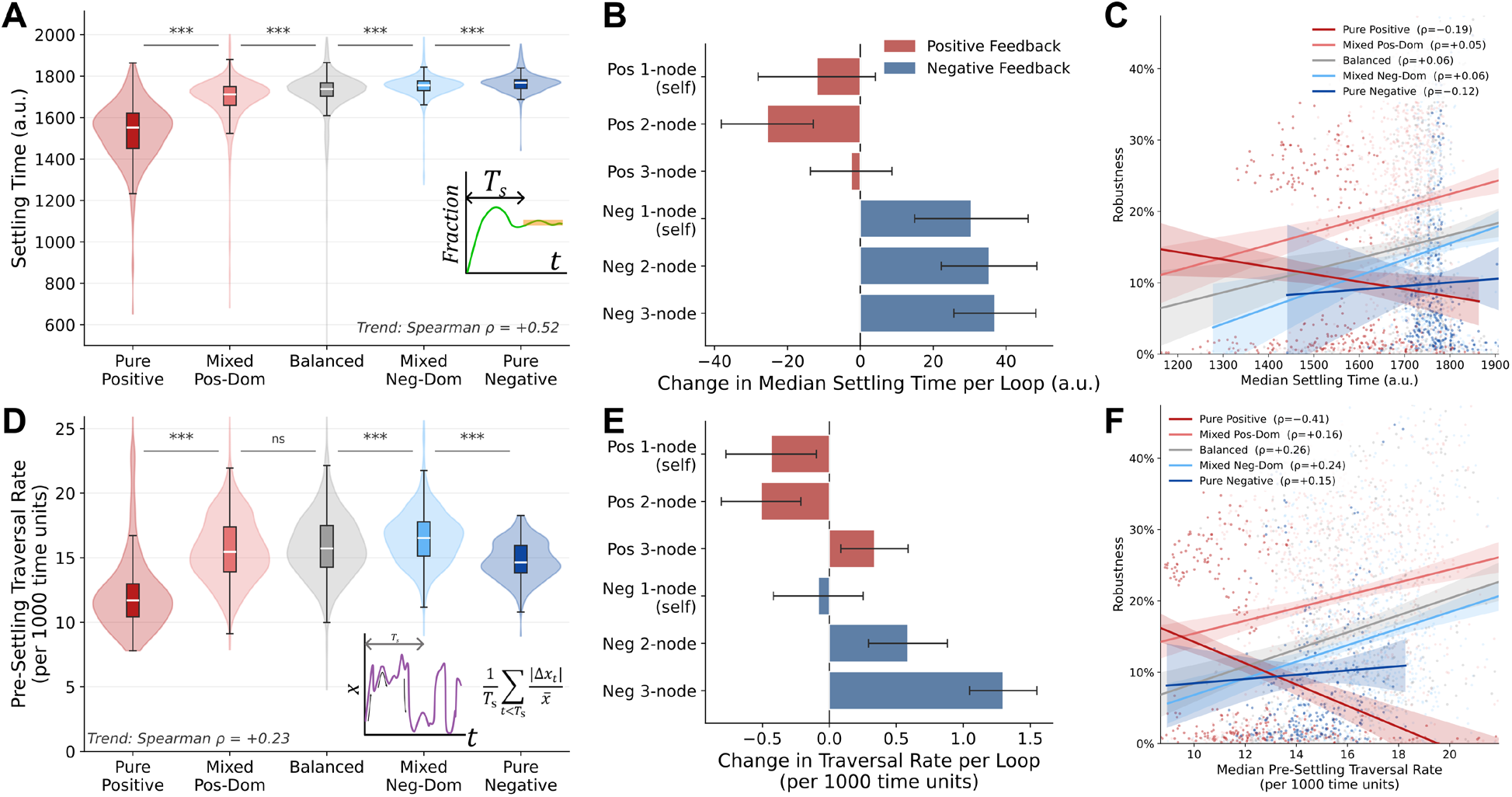
Dynamics of HR and their associations with network topology and robustness. **A**. Distributions of network-wise median settling time for 5 categories of networks. **B**. Effects of loop components (predictor variables in OLS) in terms of change in median settling time per loop. **C**. Distributions of and associations between median settling time and HR robustness. **D**. Distributions of median pre-settling traversal rate of all state variables in single cells of 5 categories of networks. **E**. Effects of loop components (predictor variables in OLS) in terms of change in median pre-settling traversal rate per loop. **F**. Distributions of and associations between median pre-settling fluctuation and HR robustness. All error bars indicate 95% confidence interval for sampling uncertainty. ns: *p* > 0.05. ***: *p* < 0.001.

HR requires individual cells to deviate from their original state and switch into a new state, and this transition is governed by alteration of gene expression levels (**Figure 2B** callout). We asked whether HR robustness is correlated with how fast gene expression changes during the state switches before HR. To quantify the single-cell-level “costs” of HR in terms of alterations of gene expression, we used a pre-settling traversal rate, defined by the sum of the absolute changes in transcription factor activity from the start of the simulation up to the parameter set’s settling time, normalized by the total settling time (see Methods). We found that positive feedback loops had the lowest traversal rates, whereas the mixed negative-dominant networks had the highest traversal rates among HR-enabling networks (**Figure 4D**). Together with the differential settling times, our results suggest that positive feedback can achieve HR quickly without requiring each cell to undergo rapid state changes in gene expression space. Conversely, negative feedback-governed HR may need oscillation, which requires gene expression to go back-and-forth between two states for multiple rounds before settling. Consistent with this hypothesis, the number of 3-node NFBLs known to generate oscillations was strongly positively associated with pre-settling traversal rate (**Figure 4E**). In addition, we found that more negative feedback-containing networks had “overshoot” dynamics of cell fractions before settling than pure positive feedback networks did (**Figure S4**). Interestingly, the pre-settling traversal rate was positively correlated with HR robustness in all categories except pure positive feedback networks, in which the increased median traversal rate reduced HR robustness (**Figure 4F**, **Figure S3 and Figure S5**).

In summary, our results showed two main differences between positive feedback and negative feedback in HR dynamics: positive feedback accelerates HR and it achieves HR through smaller changes of gene expression per unit time, whereas negative feedback slows down HR and it achieves HR through large changes of gene expression per unit time.

### The impact of noise variations on heterogeneity restoration

We next asked how the level of gene expression noise can influence HR and its dynamics. We repeated all simulations at two additional noise levels: first decreasing noise by 5-fold and then increasing it by 5-fold. We compared HR robustness and dynamics across the three noise levels, 0.2 (low), 1 (basal) and 5 (high). Interestingly, both noise reduction and amplification resulted in the decrease of the number of HR-enabling networks across all five categories of feedback composition (**Figure 5A**). High noise can destroy the bimodal pattern of gene expression in cell populations. Low noise can restrict cell state transitions required for heterogeneity recovery. In particular, for bistable (double-basin)-like systems driven by positive feedback, low noise gave rise to a dramatic reduction of HR-enabling parameter sets due to the reduced capacity of recovering heterogeneous patterns from both extreme ends of the expression spectrum (**Figure S6**), a phenomenon not observed with high noise. Although the five categories had comparable reduction in absolute percentage of HR-enabling networks upon noise variations, pure positive feedback networks had the most relative reduction. Under high noise, its disadvantage in HR existence became statistically significant compared to the mixed categories (**Figure 5A**). This result further highlights the distinction between HR and ability to produce multistability, a dynamic feature enabled by positive feedback. The alterations of noise levels in both directions also caused a decrease in HR robustness, while the mixed-feedback categories remained to be more robust than the pure feedback categories. Variations of noise reduced the HR robustness of pure positive feedback networks dramatically: the median HR robustness (the percentage of HR-enabling parameter sets per network) became less than 1% under both low and high noise conditions, making it the least robust category among all network types (**Figure 5B**). However, we observed a small subset of highly robust pure-positive-feedback networks that became prominent under high noise (**Figure 5B**, top right). We asked what network features beyond density are enriched in this network group. Strikingly, nearly 100% of the top 10% robust pure-positive-feedback networks under high noise contained mutual repression (the toggle switch motif), showing a statistically significant enrichment (**Figure 5C**). This enrichment was also observed with other network categories to the lesser extent under high noise, but only the pure positive feedback category showed this robustness-mutual-repression association across all noise levels (**Figure 5C** bottom). We found that the settling time was generally stable upon noise variations (**Figure 5D**). However, consistent with its unique negative correlation between robustness and settling time, the pure positive feedback networks had relatively longer settling times with either decrease or increase of noise (**Figure 5D**).

**Figure 5.**
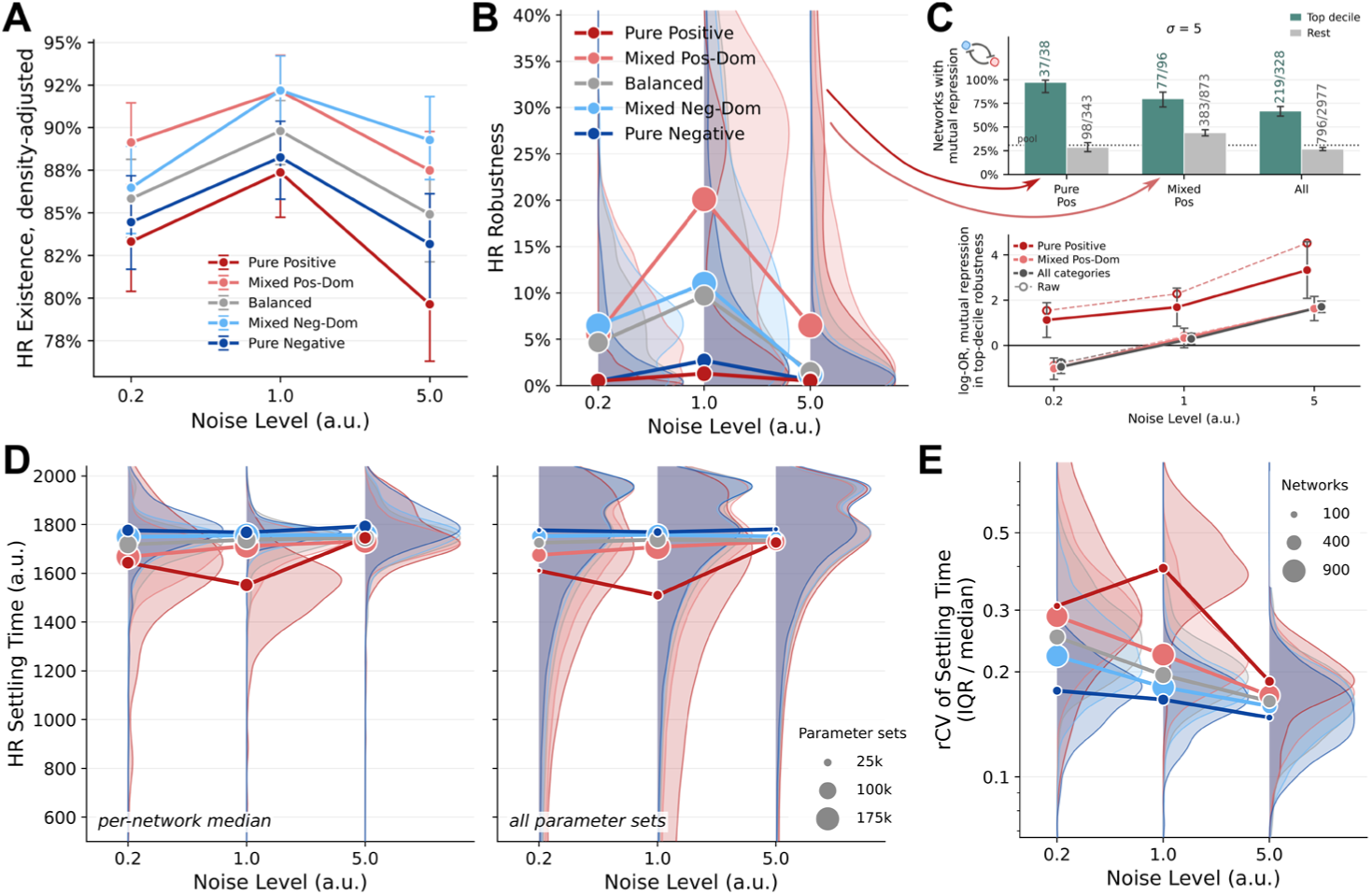
Influence of noise on HR in multiple categories of GRNs. **A**. Density-adjusted HR rate by networks under three levels of noise with g-computation from a logistic model in *n*_edge_+ *n*_edge_^2^, with 95% network-bootstrap CIs. **B**. Distributions of median HR robustness (percentage of parameter sets) for indicated categories of networks with 3 levels of noise. KDE plots show distributions of network means. Dots show means of mean robustness for each category. **C**. Top: Fractions of mutual-repression-containing networks among high robustness (top 10%) are shown in green, and those among other networks are shown in gray. Bottom: Log-odds ratios for mutual-repression-containing and high robustness under 3 levels of noise. Density correction was performed with *n*_edge_-based strata and pooled by the Mantel–Haenszel method. Error bars show 95% confidence intervals. **D**. Left: Distributions of median HR settling time for indicated categories of networks. KDE plots show distributions of network medians. Dots show means of median HR times for each category. Right: Distributions of parameter set-wise HR settling time for indicated categories of networks. KDE plots show distributions of HR settling times of all parameter sets in each category. Dots show medians of HR times for all parameter sets in each category. **E**. Summary variability quantification (interquartile range / median) for networks with more than 10 HR parameter sets.

Since our simulation data for HR settling time was collected for each parameter set, we wondered if tuning parameter values (i.e. changing kinetic rate constants) within each network would result in meaningful changes in HR settling time. Biologically this corresponds to whether the timescale of heterogeneity recovery can be tuned with either (micro-)environmental changes or evolution without structural alterations of GRNs. Parameter set-wise distributions showed that all categories of networks contained parameter sets that gave rise to broad distributions of settling times (**Figure 5C** right), but it was unclear whether these networks have high intra-network variabilities of HR settling times. We therefore examined settling times’ robust coefficients of variation (rCV) across parameter sets within individual networks (**Figure 5E**), which give a direct quantification of tunability. Interestingly, while pure positive feedback networks had the highest intra-network variations (i.e. tunability) across all noise levels, the tunability is achieved at the expense of robustness and it dramatically decreased with changes of noise levels. In contrast, mixed positive-dominant networks had both high tunability and high HR robustness across noise levels, suggesting that tunability of settling time can be an advantage of this network category without the tradeoff with HR robustness. Not surprisingly, we found that the noise level correlates with traversal rate positively but increase and decrease in noise reduced the advantage of positive feedback in minimizing traversal rate (**Figure S7**).

In summary, the results from our mathematical models provided insights into both dynamical features and mechanisms of HR-enabling networks with different types of feedback loops, and we listed these insights in **Table 2**.

**Table 2.**
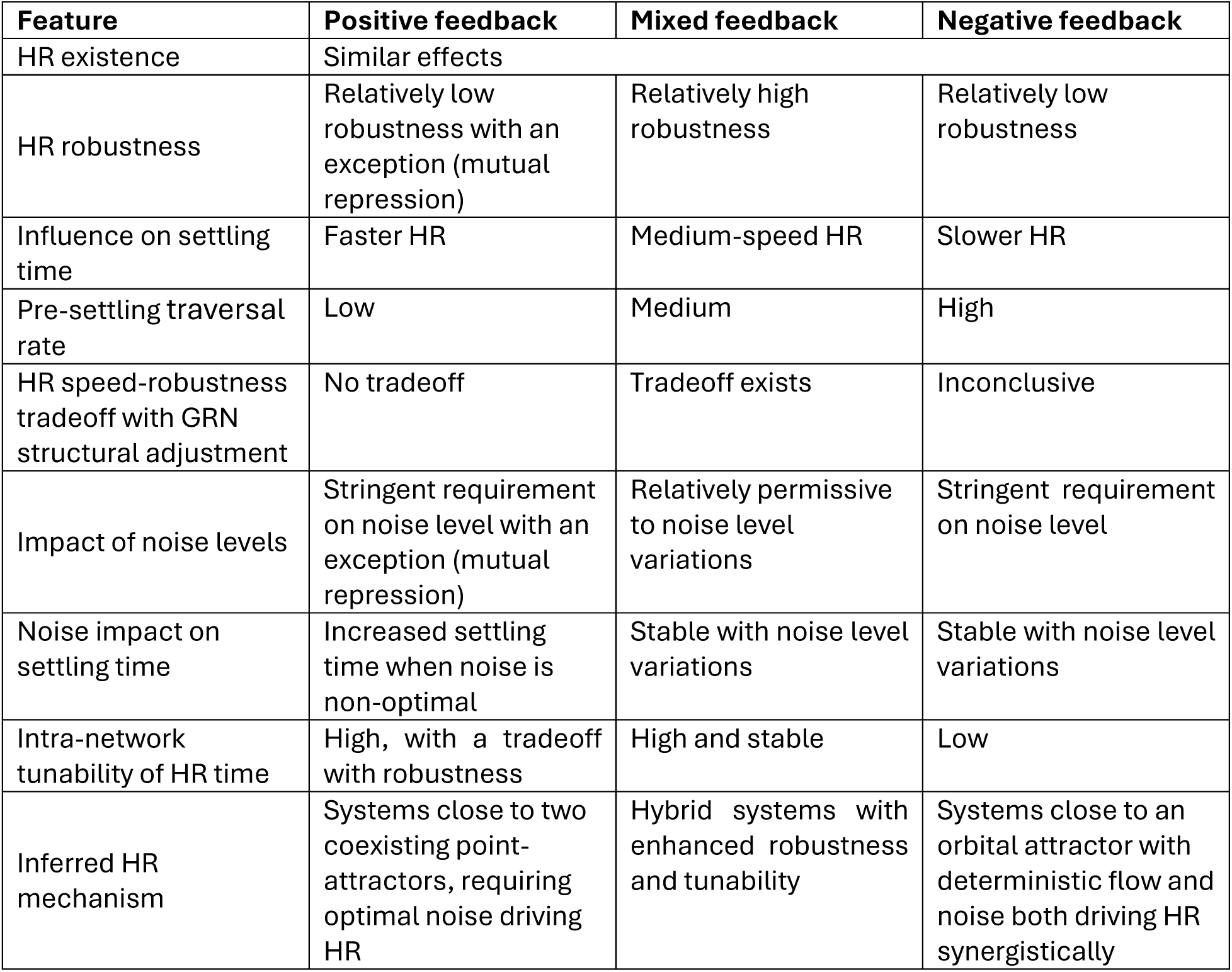
Performance and mechanism features of three types of HR-enabling networks.

| **Feature** | **Positive feedback** | **Mixed feedback** | **Negative feedback** |
|---|---|---|---|
| HR existence | Similar effects |  |  |
| HR robustness | Relatively low robustness with an exception (mutual repression) | Relatively high robustness | Relatively low robustness |
| Influence on settling time | Faster HR | Medium-speed HR | Slower HR |
| Pre-settling traversal rate | Low | Medium | High |
| HR speed-robustness tradeoff with GRN structural adjustment | No tradeoff | Tradeoff exists | Inconclusive |
| Impact of noise levels | Stringent requirement on noise level with an exception (mutual repression) | Relatively permissive to noise level variations | Stringent requirement on noise level |
| Noise impact on settling time | Increased settling time when noise is non-optimal | Stable with noise level variations | Stable with noise level variations |
| Intra-network tunability of HR time | High, with a tradeoff with robustness | High and stable | Low |
| Inferred HR mechanism | Systems close to two coexisting point-attractors, requiring optimal noise driving HR | Hybrid systems with enhanced robustness and tunability | Systems close to an orbital attractor with deterministic flow and noise both driving HR synergistically |

### Feedback-involving genes are enriched with cell fate related functions

To identify feedback-involving genes involved in experimentally validated networks, we first combined multiple sources to derive a realistic, regulator-only network suitable for topological analysis (**Figure 6A**). A network containing 4,298 regulator genes (effector-only genes were excluded because they cannot be involved in feedback loops) were obtained by pooling regulations from TRRUST, SIGNOR and RegNetwork, which contain signed, directed interactions of genes ^17–19^. Because feedback loops may appear by chance in dense networks, we used a scheme to generate a density-controlled null model by randomly choosing nodes for each edge with degree preserved degree distributions, and we identified meaningful sets of feedback-involving genes by comparing the true network with the null model (**Figure 6A** and **B**). A permissive criterion and a stringent criterion were both used to select feedback-involving genes (**Figure 6B**). To assess the functional enrichment, we generated three sets of genes related to cell fate: a single gene ontology term ‘cellular development’ (GO:0048869), a combined set from the gene ontology term ‘cell differentiation’ and ‘regulation of cell differentiation’ (GO:0030154 and GO:0045595) ^20^, and a combined set related to stem cells from MSigDB ^21–26^ (see Methods) (**Figure 6B**). We found that the enrichment of all three cell-fate-related functions in feedback-involving genes were moderate but significant (**Figure 6C**). Similar to our results showing comparable effects of positive and negative feedback in supporting the existence of HR (**Figure 3**), these two types of feedback loops did not show a significant difference in the enrichment of cell fate functions (**Figure 6C**). Although individual loop length had weak functional associations, two-node feedback loops appeared to have stronger enrichment compared to other feedback lengths (**Figure 6D**).

**Figure 6.**
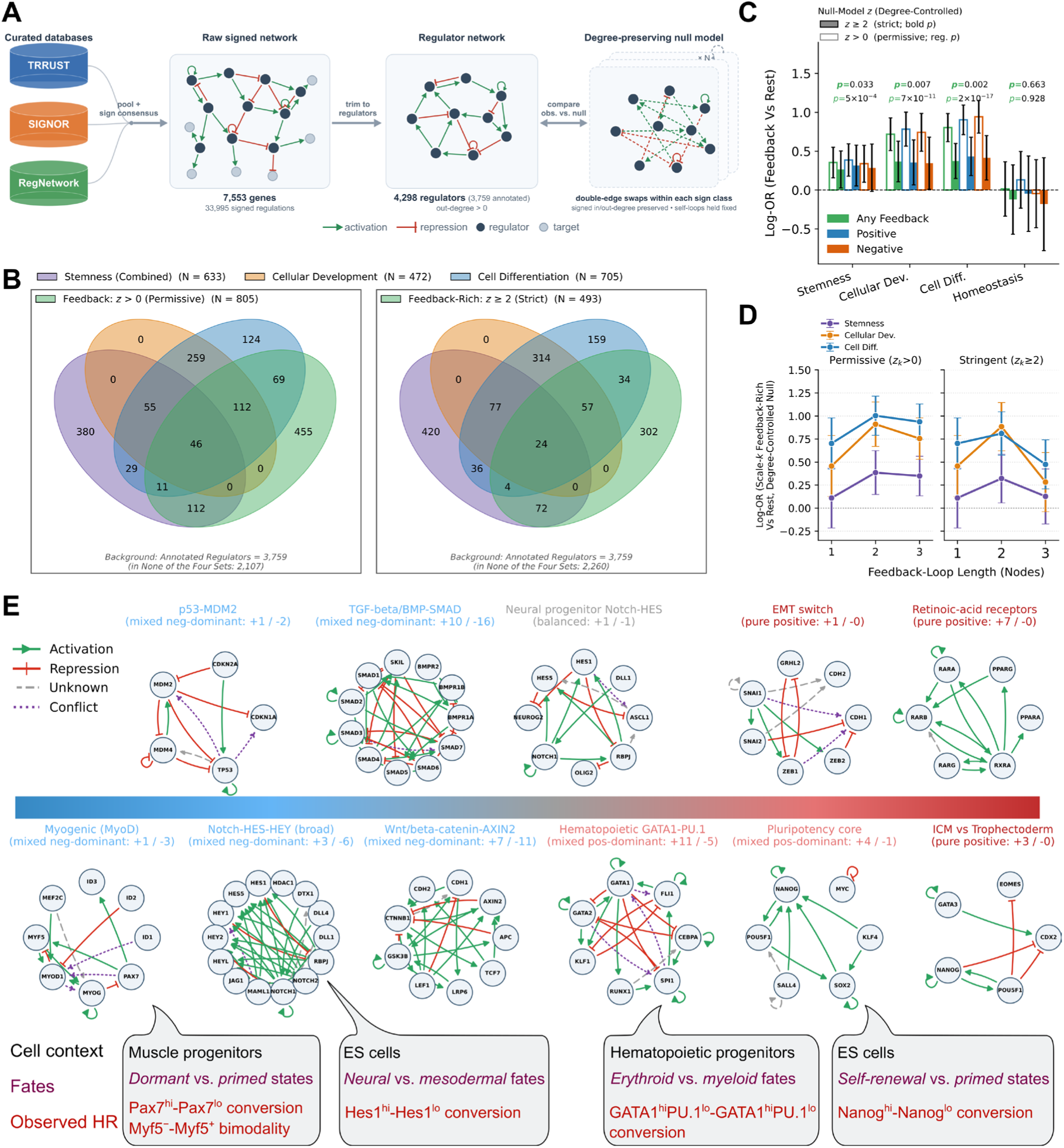
Functional enrichment of genes involved in feedback loops. **A**. Flowchart for curation of signed regulator network and generation of degree-preserving null model for quantifying feedback-involving circuits and genes. **B**. Three functional gene sets and their overlaps with feedback-involving genes (selected with two thresholds). **C**. Log-odds ratios of functional enrichments using various categories of feedback-involving genes. P values show Fisher’s exact test results for the “Any Feedback” group. A control gene set homeostasis is used for comparison. The difference between positive and negative feedback groups was not significant (*p*=0.92). **D**. Length-specific log-odds ratios compared with density-controlled null. Left and right panels show results with two thresholds for selecting feedback-involving genes. **E**. Representative GRNs controlling cell fates in specific contexts. Numbers in parentheses show the counts of positive and negative feedback loops. Conflicting edges shown in purple dashed lines indicate inconsistent regulatory signs across databases. Edges with conflicting and unknown signs are not included in feedback counts. Callouts show information about representative systems where heterogeneity recovery was observed.

We next looked for published studies on specific progenitor cells and their multi-lineage differentiation potentials and connected them to curated gene regulatory networks. Out of eleven example circuits supporting known heterogeneity recovery processes (**Figure 6E**), eight of them are with both positive and negative feedback loops, whereas the remaining three are pure positive feedback networks but they may contain negative feedback loops not included in the databases that we used (**Figure 6E**). These systems include interconvertible states in muscle progenitor cells, embryonic stem cells, hematopoietic progenitor cells and lung epithelial cells (**Supplementary Table S3**) ^1,4,27–33^. Nonetheless, we expect that there may be a wide spectrum of feedback compositions in real HR-enabling gene networks which take advantage of features of different feedback types (**Table 2**). In summary, we found that genes involved in realistic feedback loops are enriched with cell fate functions that depend on heterogeneity recovery, and we connected several experimentally observed HR systems with lineage-specific relevance to real gene circuits that may drive HR.

## Discussion

### Gene network topologies for both single-cell and tissue-level dynamics

Previous computational studies for comprehensively exploring functions of gene network topologies often focused on dynamic features that are described as either population averages or behaviors of individual cells rather than patterns across scales of genes, cells and tissues ^34–37^. In contrast, collective dynamical behaviors of populations emerging from gene regulations in single cells were often studied with either a small number of cells or a small number of networks ^38,39^. Network topologies for widely observed heterogeneity within clonal populations ^1,4,5,7,8^ had not been systematically analyzed due to limitations of computational frameworks. In this work, we performed both detailed comparisons among key network motifs and comprehensive exploration of all 3-node networks for a biological phenomenon across single-cell and population scales. Expanding the scope to modeling the collective behavior of several hundred cells across scales with a systematic search of network topologies bridges a crucial gap in this area.

Our work shows a convergent biological function of two types of circuits: positive and negative feedback loops that were previously considered to play distinct roles in biology ^40^. We found that both types of circuits can produce robust heterogeneity restoration crucial for progenitor cells’ self-renewal capabilities. This function relies on the ability of both circuit types to create attractors beyond a single point attractor through either bistability (positive feedback) or limit cycles (negative feedback), and this expansion of attractors helps to generate or restore multimodal patterns of gene expression. Nonetheless, our results show that the two types of circuits have distinct features that may be utilized by different biological systems or in different scenarios.

### Significance for lineage bias in progenitor cell populations

The heterogeneity of a progenitor cell population is associated with its differentiation potential towards different lineages. For example, it was shown that the differentiation of hematopoietic stem cells into megakaryocytic–erythroid and granulocytic–monocytic lineages can be biased in distinct subpopulations within these stem cells, whereas the balance of the differentiation potentials is accompanied by the heterogeneity restoration of gene expression in the stem cell population ^1^. This remarkable phenomenon led to the hypothesis that gene regulatory circuits such as the toggle switch between GATA1 and PU.1, lineage-determining, mutually repressive transcription factors for committing to two lineages, can govern this differentiation potential bias in hematopoietic stem cells ^41,42^. However, later studies rejected this hypothesis with findings showing a poor correlation between GATA1-PU.1 fluctuations and differentiation potentials ^43^. Our results show that while a mutual inhibition circuit is one plausible network topology for heterogeneity restoration, other gene network architectures, including those involving negative feedback, can be an effective alternative mechanism. Such a mechanism can explain the uncoupling between the bias of the initial transcription factor expression and the ultimate lineage-determining events during differentiation. Nonetheless, future work is still needed to identify specific HR-enabling circuits responsible for dynamics in progenitor cells such as hematopoietic stem cells.

### The role of noise

Fluctuations of gene expression are a type of noise that can be a source of perturbations potentially harmful for robust developmental programs. Mechanisms for noise reduction have therefore been investigated in previous theoretical and experimental studies ^35,37,44–47^. However, noise can play beneficial roles in some biological systems in a variety of ways, including sharpening tissue boundaries during development ^9,48–50^. Previous work showed that noise can change the landscape of multistable systems for cell lineage transitions ^51^. In this work, we showed that an optimal level of noise can synergize with either toggle switches or oscillators to facilitate heterogeneity restoration. While this is unsurprising, we found that noise plays different roles when working with these two types of circuits. Positive feedback-driven heterogeneity (related to bistability) is extremely sensitive to noise, which is essential for cells to move between basins of attractions. While noise can desynchronize single-cell oscillations in the population to achieve heterogeneity, in this scenario the restoration performance is less sensitive to the noise level. Our work revealed the differential importance of noise when combined with different types of gene circuits.

### Heterogeneity restoration on different timescales

Our results showed that heterogeneity restoration can be achieved with speeds that can vary significantly both across different network topologies and across different parameter sets for the same network topology. In addition to cell-autonomous control of the heterogeneity restoration speed, a recent study showed that cell-cell communications can influence the speed of rebalancing subpopulations of cancer cells ^52^. While in our analysis we assumed that fast restoration is a performance advantage for simplicity, slow restoration to balanced subpopulations can result in stability of gene expression and lineage commitment potential at the single-cell level. We expect that some progenitor cell populations can use a combination of fast and slow restoration mechanisms to further diversify the strategies and dynamical features that allow optimal self-renewal. With experiments in a limited time window, it was previously reported that the subpopulations in hematopoietic stem cells responsible for self-renewal are different from those with biased differentiation potentials ^2^. Our results suggest that the attractors of these two groups of cells, whether they are within a single epigenetic landscape or different ones, may be shaped by two types of gene regulatory networks. Future work is warranted to identify how a clonal population can produce such distinct attractor types governed by different subnetworks of the gene regulation systems. Our work provides a comprehensive mapping of gene regulatory circuits to heterogeneity restoration at both single-cell and population levels, and a foundation for future identification of specific gene circuits that can govern self-renewal at various timescales.

## Methods

### Network selections

We systematically generated structurally unique topologies for one, two or three genes (1-3 nodes) by creating a collection of 3×3 adjacency matrices where each interaction weight is from the set (−1, 0, 1). An activation interaction is represented by +1, an inhibitory interaction is denoted by −1, and 0 denotes the absence of a connection between two genes. We started with 3^9^ =19,683 possible combinations of edges, and we eliminated isomorphic networks by enumerating all potential node permutations. We also removed disjoint networks (their network topologies are captured by smaller networks) and the null network. The final number of distinct networks is 3325, each of which represents a unique functional topology.

### Mathematical models

We used ordinary differential equations (ODEs) to model systems containing one to three genes (1-3 nodes) for the 3325 selected networks. Each ODE describes the activity dynamics of a lineage-determining transcription factor (TF), and it has the dimensionless form

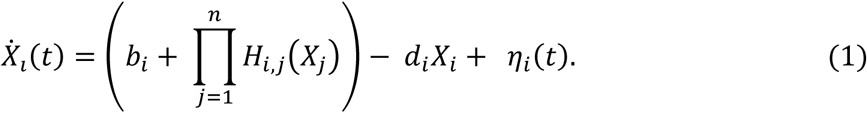

Here, X*_i_* ( *i* ∈ (1,2,3)) represents the activity of a transcription factor (TF). *b_i_* is the basal production rate of the TF in the absence of any regulator. The terms inside the parentheses represent the production rate under the control of transcriptional activators and inhibitors. The regulation is determined by the product of Hill functions (*H_i,j_*), representing synergistic or "AND-gate" logic. For each regulating TF (X*_j_*), the Hill function is defined as

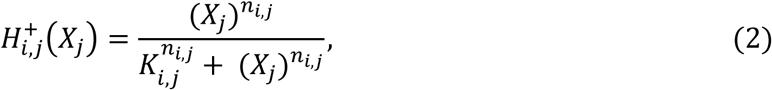

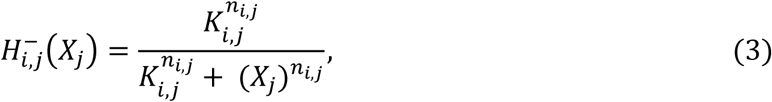

depending on whether the regulator is an activator or an inhibitor. *K_i,j_* is the apparent dissociation constant of the regulating TF binding to its regulatory element of the promoter, and it describes the effectiveness of the regulation in terms of the concentration of the TF. *n_i,j_* is the Hill coefficient. *d_i_* is the degradation rate constant. The production rate of the proteins is assumed to be linearly correlated with the mRNA production rate. We modeled noise *η_i_*(*t*) as an Ornstein-Uhlenbeck (OU) process to describe the temporal persistence of biological fluctuations. This represents extrinsic noise with a correlation time *τ* and magnitude *σ_i_*, governed by

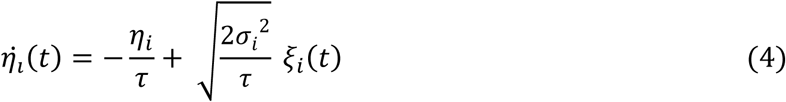

where *ξ_i_*(*t*) describes standard Gaussian white noise. When a deterministic version of the model is needed to determine a fixed point, we set the noise terms to zero. Here, all the parameters are dimensionless, and one time unit of our model corresponds to 9 min. The OU correlation time was fixed at *τ* = 0.1. Kinetic parameters (*b_i_*, *d_i_*, *K_i,j_*, *n_i,j_*) were drawn from truncated log-normal distributions bounded by *b_i_* ∈ [0.001, 0.1], *d_i_* ∈ [0.01, 0.5], *K_i,j_* ∈ [0.1, 5.0], and *n_i,j_* ∈ [1, 10]. We repeated the full pipeline at three noise magnitudes, *σ_i_* ∈ {0.2, 1.0, 5.0} (*σ_i_* = 1.0 as basal), applied to all three TFs.

To describe the behavior of the cell population, we solved the stochastic ODEs (SODEs) *n* times with 2000 time-units, where *n* represents the number of cells. Each trajectory represents a trajectory of one cell. We utilized a total of *n* = 500 trajectories for each individual network configuration across 1,000 randomly generated parameter sets from truncated log-normal distributions. To select initial conditions, we performed a deterministic pre-run for each parameter set by solving the ODE model over the interval t ∈ [0, 2000]. We then extracted the X*_i_* state level at t=2000 (either at a fixed point or a stable orbit) and used these values to define the initial conditions for the subsequent stochastic simulations. This method ensures that all SODE trajectories originate from a post-transient, biologically stable state. The distribution at the stationary phase (or the final time point) represents the long-term behavior of the population and its heterogeneity is quantified with the procedure described below in the next subsection.

To assess whether the identified phenotypic states were robust to the choice of initial condition (i.e. HR is possible from each of the modes of the distribution), rather than contingent on the basal pre-run state, we conducted a second series of stochastic simulations using the parameter sets previously classified as multimodality-enabling. The mean activity level of the alternative phenotypic subpopulation, defined as the cluster whose average state was more distant from the original deterministic initial condition, was determined from the results of the original simulation series and subsequently used to initialize new stochastic trajectories, replacing the basal pre-run state as the initial condition. This alternative-initial-condition simulation series was conducted at each of the three noise magnitudes examined as a filtering step.

### Heterogeneity recovery

The definition of heterogeneity recovery (HR) is based on the ability for a cell population with a uniform initial condition to achieve multimodal gene expression at the stationary phase (multimodality). This ensures that the heterogeneity that we study here contains separable subpopulations with cells poised to different lineage commitments but still capable of multipotency renewal. We require HR to be bidirectional, i.e. recovery should be initiated with each mode of the stationary distribution. We investigated the distribution of TF activities at the final time point (t = 2000) to determine multimodality within the simulated cell population. We used Gaussian Mixture Models (GMM) to assess whether the terminal-state distribution of each parameter set had resolved into distinct phenotypic subpopulations, an approach previously applied to recover heterogeneity from experimental data ^1^. For each candidate distribution, we compared a single-component and a two-component mixture model and used the Bayesian Information Criterion (BIC) to select between them: a distribution was considered a candidate for multimodality only if the two-component model achieved a lower BIC than the single-component model.

We used two additional criteria to select HR-enabling parameters. First, the two subpopulations must be clearly separated: their means must differ by at least 30% from the larger one along every dimension (TF) used for GMM models. Second, the two subpopulations had to be reasonably balanced in size: each must contain at least 10% of the simulated cells. A parameter set was classified as meaningfully heterogeneous, or HR-enabling, only if the two-component model was preferred by BIC and met all these conditions. Otherwise, it was classified as non-HR-enabling. For networks with one active transcription factor, we fit the model to the distribution of that factor’s activity alone. For networks with two or more TFs, we fit a two-factor model to each possible pair of TFs. A parameter set was retained only if all pairwise fits satisfied the BIC and separation criteria described above.

### Metrics for HR dynamics

The existence of HR for a network is defined by whether any of its 1,000 parameter sets produced HR, i.e. bidirectional multimodality regeneration. Similarly, the robustness of HR is defined as the percentage of HR-enabling parameter sets for a network. For simulations from an HR-enabling parameter set, the steady state value of a type of cell is defined as the mean value of its fraction over the final 200 time units of the simulation. The settling time was then defined as the first timepoint after which the cell fraction stays within a 5% (in absolute cell fraction) band from its steady-state value.

For the time-course fraction of the non-starting cell type, we checked whether it reached a peak value more than 10% higher than its final value. When such a peak value was found, we recorded the existence of overshoot for this parameter set as a metric for fluctuating cell fractions before settling. To measure the extent to which individual cells alter their expression before settling into their final phenotype (this can be viewed as the single-cell “cost” for HR), we calculated a pre-settling traversal rate for each cell trajectory: the sum of the absolute changes in transcription factor activity from the start of the simulation up to the parameter set’s settling time normalized by the total settling time, i.e.

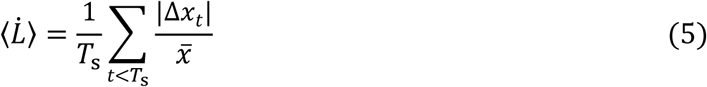

where *T*_s_ is the settling time and *x̅* is the mean expression level of a TF. The mean pre-settling traversal rate across all modeled TFs was used to describe a per-set property, whereas the median pre-settling traversal rate across all parameter sets for an HR-enabling network was used to give a representative value for the network.

### Statistical analysis and predictive models

We used several schemes to categorize networks for our statistical analyses: 1) a basic binary categorization of whether each network has any feedback or feed-forward loops; 2) an elementary feedback composition (positive/negative feedback, 1-node/2-node/3-node loops); 3) the ratio between positive and negative feedback loops (purely positive, mixed positive-dominant, balanced, mixed-negative-dominant and purely negative); 4) feedforward-loop coherence; and 5) density (number of edges). Associations between a binary topology feature and a categorical outcome are quantified as log odds ratios from 2×2 tables with a Haldane–Anscombe correction, and Fisher’s exact test for the raw *p*-value. Because feedback categories differ in density, contribution is also reported density-adjusted by one of two routes. First, Mantel–Haenszel pooling over strata of number of edges, with the Cochran–Mantel–Haenszel test and a Breslow–Day test of odds-ratio homogeneity. Secondly, a binomial GLM on density with g-computation over the observed density distribution and percentile intervals from a network-level bootstrap (circuits are resampled). We used per-loop effects on rate outcomes to build a binomial GLM giving log-odds per additional loop with quasi-binomial intervals, whereas per-loop effects on continuous outcomes were from OLS with HC3 robust standard errors, fitted without and with the number of edges included. Spearman correlations are density-controlled by partial Spearman on number of edges. Group comparisons were performed with Kruskal–Wallis and Mann–Whitney U tests with Cliff’s delta and Holm-corrected *p*-values for adjacent-category contrasts, the Wilcoxon signed-rank test for paired within-circuit comparisons, and the Friedman test for repeated measures across noise. Monotone trends were quantified with Spearman rank correlation coefficient in the ordinal category. We used effect sizes rather than *p*-values for interpretation. To quantify predictability, we used random-forest and gradient-boosting models under repeated (15 times) or nested 5-fold cross-validation, scored by R² and Spearman correlation coefficients for continuous labels and ROC-AUC plus minority-class PR-AUC for HR existence, each against a null model with randomized labels.

### Validated gene circuits and functional enrichment analysis

Three databases containing signed, directed gene regulations, TRRUST v2, SIGNOR and RegNetwork ^17–19^, were pooled across human and mouse, with mouse genes mapped to human orthologs and duplicate edges merged into a consensus sign. Within a feedback-capable subnetwork, every simple directed loop of one to four genes was enumerated and labeled by size and by the feedback sign, which gave each gene a feedback profile. Functional gene sets were obtained from GO Biological Process and expanded to descendants ^20^, with the stem/progenitor set supplemented from MSigDB ^21–26^. Enrichment was first measured as a log-odds ratio with Fisher exact tests using Mantel–Haenszel degree-quintile matching and logistic regression on log-degree. Next, we used a degree-preserving edge-swap null model (200 randomizations) to convert each gene’s loop count into a z-score. Bonferroni-corrected results were used at various levels of the GO hierarchy to localize the GO-based enrichment.

## Author contributions

Suchana Chakravarty: Methodology, Investigation, Writing—original draft, Writing—review C editing. Rija Tonny Christian Ramarolahy: Methodology, Writing—review C editing. Tian Hong: Project administration, Supervision, Resources, Methodology, Writing—original draft, Writing—review C editing.

## Data availability

Computer code for reproducing results in this manuscript is available at the GitHub repository https://github.com/SuchanaChakravarty/GRN_HR.

## AI Statement

AI tools have been used for organizing computer code and for correcting grammatical and typographical errors in the manuscript.

## References

1. Chang, H.H., Hemberg, M., Barahona, M., Ingber, D.E., and Huang, S. (2008). Transcriptome-wide noise controls lineage choice in mammalian progenitor cells. Nature 453, 544–547. 10.1038/nature06965.

2. Pina, C., Fugazza, C., Tipping, A.J., Brown, J., Soneji, S., Teles, J., Peterson, C., and Enver, T. (2012). Inferring rules of lineage commitment in haematopoiesis. Nat. Cell Biol. 14, 287–294.

3. Strebinger, D., Deluz, C., Friman, E.T., Govindan, S., Alber, A.B., and Suter, D.M. (2019). Endogenous fluctuations of OCT4 and SOX2 bias pluripotent cell fate decisions. Mol. Syst. Biol. 15, MSB199002.

4. Chakraborty, M., Hu, S., Visness, E., Del Giudice, M., De Martino, A., Bosia, C., Sharp, P.A., and Garg, S. (2020). MicroRNAs organize intrinsic variation into stem cell states. Proc. Natl. Acad. Sci. USA 117, 6942–6950.

5. Kalmar, T., Lim, C., Hayward, P., Muñoz-Descalzo, S., Nichols, J., Garcia-Ojalvo, J., and Martinez Arias, A. (2009). Regulated fluctuations in nanog expression mediate cell fate decisions in embryonic stem cells. PLoS Biol. 7, e1000149.

6. Jordan, N.V., Bardia, A., Wittner, B.S., Benes, C., Ligorio, M., Zheng, Y., Yu, M., Sundaresan, T.K., Licausi, J.A., and Desai, R. (2016). HER2 expression identifies dynamic functional states within circulating breast cancer cells. Nature 537, 102–106.

7. Spencer, S.L., Gaudet, S., Albeck, J.G., Burke, J.M., and Sorger, P.K. (2009). Non-genetic origins of cell-to-cell variability in TRAIL-induced apoptosis. Nature 459, 428–432.

8. Singh, D.K., Ku, C.J., Wichaidit, C., Steininger, R.J., Wu, L.F., and Altschuler, S.J. (2010). Patterns of basal signaling heterogeneity can distinguish cellular populations with different drug sensitivities. Mol. Syst. Biol. 6, MSB201022.

9. Huang, S. (2009). Non-genetic heterogeneity of cells in development: more than just noise. Development 136, 3853–3862.

10. Yu, P., Nie, Q., Tang, C., and Zhang, L. (2018). Nanog induced intermediate state in regulating stem cell differentiation and reprogramming. BMC Syst. Biol. 12, 22.

11. Lahmann, I., Zhang, Y., Baum, K., Wolf, J., and Birchmeier, C. (2021). An oscillatory network controlling self-renewal of skeletal muscle stem cells. Exp. Cell Res. 409, 112933.

12. Farjami, S., Camargo Sosa, K., Dawes, J.H.P., Kelsh, R.N., and Rocco, A. (2021). Novel generic models for differentiating stem cells reveal oscillatory mechanisms. Journal of The Royal Society Interface 18, 20210442.

13. Phillips, N.E., Manning, C.S., Pettini, T., Biga, V., Marinopoulou, E., Stanley, P., Boyd, J., Bagnall, J., Paszek, P., and Spiller, D.G. (2016). Stochasticity in the miR-9/Hes1 oscillatory network can account for clonal heterogeneity in the timing of differentiation. eLife 5, e16118.

14. Kobayashi, T., and Kageyama, R. (2011). Hes1 oscillations contribute to heterogeneous differentiation responses in embryonic stem cells. Genes 2, 219–228.

15. Clauss, B., and Lu, M. (2023). A quantitative evaluation of topological motifs and their coupling in gene circuit state distributions. Iscience 26.

16. Huang, L., Clauss, B., and Lu, M. (2022). What makes a functional gene regulatory network? A circuit motif analysis. The journal of physical chemistry. B 126, 10374.

17. Han, H., Cho, J.-W., Lee, S., Yun, A., Kim, H., Bae, D., Yang, S., Kim, C.Y., Lee, M., and Kim, E. (2018). TRRUST v2: an expanded reference database of human and mouse transcriptional regulatory interactions. Nucleic Acids Res. 46, D380–D386.

18. Lo Surdo, P., Iannuccelli, M., Karis, K., Meo, E., Omidi, P., Tosoni, M., Graziosi, S., Panni, S., Fiorillo, M.T., and Licata, L. (2026). SIGNOR 4.0: the 2025 update with focus on phosphorylation data. Nucleic Acids Res. 54, D682–D690.

19. Liu, Z.-P., Wu, C., Miao, H., and Wu, H. (2015). RegNetwork: an integrated database of transcriptional and post-transcriptional regulatory networks in human and mouse. Database 2015, bav095.

20. Carbon, S., Douglass, E., Good, B.M., Unni, D.R., Harris, N.L., Mungall, C.J., Basu, S., Chisholm, R.L., Dodson, R.J., and Hartline, E. (2021). The Gene Ontology resource: enriching a GOld mine. Nucleic Acids Res. 49, D325–D334.

21. Ben-Porath, I., Thomson, M.W., Carey, V.J., Ge, R., Bell, G.W., Regev, A., and Weinberg, R.A. (2008). An embryonic stem cell-like gene expression signature in poorly differentiated aggressive human tumors. Nat. Genet. 40, 499–507. 10.1038/ng.127.

22. Bhattacharya, B., Miura, T., Brandenberger, R., Mejido, J., Luo, Y., Yang, A.X., Joshi, B.H., Ginis, I., Thies, R.S., Amit, M., et al. (2004). Gene expression in human embryonic stem cell lines: unique molecular signature. Blood 103, 2956–2964. 10.1182/blood-2003-09-3314.

23. Müller, F.-J., Laurent, L.C., Kostka, D., Ulitsky, I., Williams, R., Lu, C., Park, I.-H., Rao, M.S., Shamir, R., Schwartz, P.H., et al. (2008). Regulatory networks define phenotypic classes of human stem cell lines. Nature 455, 401–405. 10.1038/nature07213.

24. Ramalho-Santos, M., Yoon, S., Matsuzaki, Y., Mulligan, R.C., and Melton, D.A. (2002). "Stemness": transcriptional profiling of embryonic and adult stem cells. Science 298, 597–600. 10.1126/science.1072530.

25. Wong, D.J., Liu, H., Ridky, T.W., Cassarino, D., Segal, E., and Chang, H.Y. (2008). Module map of stem cell genes guides creation of epithelial cancer stem cells. Cell Stem Cell 2, 333–344. 10.1016/j.stem.2008.02.009.

26. Liberzon, A., Subramanian, A., Pinchback, R., Thorvaldsdóttir, H., Tamayo, P., and Mesirov, J.P. (2011). Molecular signatures database (MSigDB) 3.0. Bioinformatics 27, 1739–1740.

27. Chazaud, C., Yamanaka, Y., Pawson, T., and Rossant, J. (2006). Early Lineage Segregation between Epiblast and Primitive Endoderm in Mouse Blastocysts through the Grb2-MAPK Pathway. Dev. Cell 10, 615–624. 10.1016/j.devcel.2006.02.020.

28. De Micheli, A.J., Spector, J.A., Elemento, O., and Cosgrove, B.D. (2020). A reference single-cell transcriptomic atlas of human skeletal muscle tissue reveals bifurcated muscle stem cell populations. Skeletal Muscle 10, 19. 10.1186/s13395-020-00236-3.

29. Kobayashi, T., Mizuno, H., Imayoshi, I., Furusawa, C., Shirahige, K., and Kageyama, R. (2009). The cyclic gene Hes1 contributes to diverse differentiation responses of embryonic stem cells. Genes Dev. 23, 1870–1875. 10.1101/gad.1823109.

30. Kuang, S., Kuroda, K., Le Grand, F., and Rudnicki, M.A. (2007). Asymmetric Self-Renewal and Commitment of Satellite Stem Cells in Muscle. Cell 129, 999–1010. 10.1016/j.cell.2007.03.044.

31. Osorno, R., and Chambers, I. (2011). Transcription factor heterogeneity and epiblast pluripotency. Philosophical Transactions of the Royal Society B: Biological Sciences 366, 2230–2237. 10.1098/rstb.2011.0043.

32. Rocheteau, P., Gayraud-Morel, B., Siegl-Cachedenier, I., Blasco, Maria A., and Tajbakhsh, S. (2012). A Subpopulation of Adult Skeletal Muscle Stem Cells Retains All Template DNA Strands after Cell Division. Cell 148, 112–125. 10.1016/j.cell.2011.11.049.

33. Bhatia, S., Gunter, J.H., Burgess, J.T., Adams, M.N., O’Byrne, K., Thompson, E.W., and Duijf, P.H.G. (2023). Stochastic epithelial-mesenchymal transitions diversify non-cancerous lung cell behaviours. Transl. Oncol. 37, 101760.

34. Ma, W., Trusina, A., El-Samad, H., Lim, W.A., and Tang, C. (2009). Defining network topologies that can achieve biochemical adaptation. Cell 138, 760–773.

35. Qiao, L., Zhao, W., Tang, C., Nie, Q., and Zhang, L. (2019). Network topologies that can achieve dual function of adaptation and noise attenuation. Cell systems 9, 271–285.

36. Ye, Y., Kang, X., Bailey, J., Li, C., and Hong, T. (2019). An enriched network motif family regulates multistep cell fate transitions with restricted reversibility. PLoS computational biology 15, e1006855. 10.1371/journal.pcbi.1006855.

37. Qiao, L., Zhang, Z.-B., Zhao, W., Wei, P., and Zhang, L. (2022). Network design principle for robust oscillatory behaviors with respect to biological noise. Elife 11, e76188.

38. Jiménez, A., Cotterell, J., Munteanu, A., and Sharpe, J. (2017). A spectrum of modularity in multi-functional gene circuits. Mol. Syst. Biol. 13, 925.

39. Perez-Carrasco, R., Barnes, C.P., Schaerli, Y., Isalan, M., Briscoe, J., and Page, K.M. (2018). Combining a toggle switch and a repressilator within the AC-DC circuit generates distinct dynamical behaviors. Cell systems 6, 521–530.

40. Tyson, J.J., Chen, K.C., and Novak, B. (2003). Sniffers, buzzers, toggles and blinkers: dynamics of regulatory and signaling pathways in the cell. Curr. Opin. Cell Biol. 15, 221–231.

41. Graf, T., and Enver, T. (2009). Forcing cells to change lineages. Nature 462, 587.

42. Orkin, S.H., and Zon, L.I. (2008). Hematopoiesis: an evolving paradigm for stem cell biology. Cell 132, 631–644.

43. Hoppe, P.S., Schwarzfischer, M., Loeffler, D., Kokkaliaris, K.D., Hilsenbeck, O., Moritz, N., Endele, M., Filipczyk, A., Gambardella, A., and Ahmed, N. (2016). Early myeloid lineage choice is not initiated by random PU. 1 to GATA1 protein ratios. Nature 535, 299–302.

44. Li, C.-J., Hong, T., Tung, Y.-T., Yen, Y.-P., Hsu, H.-C., Lu, Y.-L., Chang, M., Nie, Q., and Chen, J.-A. (2017). MicroRNA filters Hox temporal transcription noise to confer boundary formation in the spinal cord. Nat. Commun. 8, 14685. 10.1038/ncomms14685.

45. Ramos, A.F., Hornos, J.E.M., and Reinitz, J. (2015). Gene regulation and noise reduction by coupling of stochastic processes. Physical Review E S1, 020701.

46. Schmiedel, J.M., Klemm, S.L., Zheng, Y., Sahay, A., Blüthgen, N., Marks, D.S., and van Oudenaarden, A. (2015). MicroRNA control of protein expression noise. Science 348, 128–132.

47. Osella, M., Bosia, C., Corá, D., and Caselle, M. (2011). The role of incoherent microRNA-mediated feedforward loops in noise buffering. PLoS computational biology 7, e1001101.

48. Holmes, W.R., de Mochel, N.S.R., Wang, Q., Du, H., Peng, T., Chiang, M., Cinquin, O., Cho, K., and Nie, Q. (2017). Gene expression noise enhances robust organization of the early mammalian blastocyst. PLoS computational biology 13, e1005320.

49. Kittisopikul, M., and Süel, G.M. (2010). Biological role of noise encoded in a genetic network motif. Proc. Natl. Acad. Sci. USA 107, 13300. 10.1073/pnas.1003975107.

50. Zhang, L., Radtke, K., Zheng, L., Cai, A.Q., Schilling, T.F., and Nie, Q. (2012). Noise drives sharpening of gene expression boundaries in the zebrafish hindbrain. Mol. Syst. Biol. 8, 613. 10.1038/msb.2012.45.

51. Coomer, M.A., Ham, L., and Stumpf, M.P.H. (2021). Noise distorts the epigenetic landscape and shapes cell-fate decisions. Cell Systems.

52. Lopez, D., Tyson, D.R., and Hong, T. (2025). Intercellular signaling reinforces single-cell level phenotypic transitions and facilitates robust re-equilibrium of heterogeneous cancer cell populations. Cell Communication and Signaling 23, 386.

