## Supplementary Information for "Gene network topologies for heterogeneity restoration during self-renewal"

---

### Estimation of parameters in mathematical models and heterogeneity recovery time

The model includes three species ($X_i, where\ i = 1,2,3$) and a 3x3 regulatory interaction matrix, resulting in a total of 24 kinetic parameters: three basal production rates ($b_i$), three first-order degradation rates ($d_i$), and two sets of nine Hill function parameters (Hill threshold constants ($K_{i,j}$) and Hill cooperativity exponents ($n_{i,j}$)), one for each entry in the regulatory matrix. We sampled each parameter from a truncated lognormal distribution from a range that we describe later. This is a typical choice for strictly positive biological variables because biological rate constants are always positive and vary over orders of magnitude [1]. We used the inverse cumulative distribution function method for sampling [2], where the log-space mean was set to the geometric midpoint of the range and the width was set so the bounds span roughly four standard deviations.

Here, the state variables are expressed as normalized dimensionless concentrations with a reference value of 1. The cooperativity exponents $n_{ij}$ are dimensionless integers. The Hill threshold constants $K_{ij}$ are expressed in the same normalized concentration units as the state variables, with the reference concentration set to 1 in model units. The basal production rates $b_i$ and degradation rates $d_i$ carry units of [a.u.$^{-1}$]. We assumed that 1 a.u. ≈ 9 minutes. The basal production rate is expressed as [time$^{-1}$], which represents the rate of growth of the normalized expression level caused by intrinsic leaky transcription. The half-lives of mammalian proteins range from 12 minutes to 10 hours. This range includes genes under active regulation. At the fast end, Holder et al. directly evaluated turnover of major cell-cycle regulators at the metaphase-anaphase transition and discovered half-lives of roughly 5 minutes for cyclin B, securin, and geminin [3], placing their degradation rates near to our top limit. Even our slowest-degrading genes are actively turned over in comparison to typical cellular proteins; at the slow end, a half-life of 10 hours indicates faster-than-average protein turnover compared to the genome-wide median of roughly 46 hours measured by Schwanhäusser et al. across more than 5000 mammalian proteins [4]. Degradation rates ($d_i$) were therefore sampled from [0.01, 0.5] a.u.$^{-1}$, equating to [0.00111, 0.0556] min$^{-1}$ in real units. The basal production rates ($b_i$) were collected from [0.001, 0.1] a.u.$^{-1}$, which corresponds to [0.000111, 0.0111] min$^{-1}$ following conversion. This estimation is to ensure that the scaled concentrations are simulated in a range suitable for numerical calculations.

Hill cooperativity exponents ($n_{i,j}$) were sampled from [1.0, 10.0] and rounded up to the nearest integer using the ceiling function, giving integer values 1 through 10. This includes non-cooperative monomer binding (n = 1) and dimerization (n = 2), as utilized in the repressilator [5] and toggle switch [6], tetrameric complexes (n = 3-4), and multisite phosphorylation systems. Nash et al. discovered that the CDK inhibitor Sic1 requires 6 of its 9 phosphorylation sites to be occupied before degradation occurs [7], resulting in high apparent cooperativity, and Goldbeter and Koshland mathematically demonstrated that cascading covalent modifications can produce apparent Hill coefficients well above four [8]. Ranges for all parameters are listed in Supplementary Table S1.

**Supplementary Table S1.** Ranges of parameter values for sampling of 3-node networks.

| Parameter | Description | Range/Value used for simulation |
|---|---|---|
| $b_i$ | basal production rate | [0.001, 0.1] |
| $d_i$ | degradation rates | [0.01, 0.5] |
| $K_{i,j}$ | Hill threshold constants of i by j, where i = j & i ≠ j | [0.1, 5.0] |
| $n_{i,j}$ | Hill cooperativity exponents of i by j, where i = j & i ≠ j | [1.0, 10.0] |

All simulations were done for 2000 time units. Chang et al. sorted mouse hematopoietic progenitor cells into discrete Sca-1 expression subpopulations and found that more than 9 days elapsed before the three fractions regenerated Sca-1 histograms similar to that of the parental population [9]. In our simulations, parameter configurations with modest switching dynamics achieve a stable population distribution when stability time > 1500 a.u. Using the integer conversion factor derived above (1 a.u. ≈ 9 minutes), this threshold corresponds to a typical predicted recovery time of

$$1500\ a.u.\ \times 9\ \min a.u.^{-1} = 13{,}500\ min \approx 9.4\ days\ \approx 9\ days\ 9\ hours$$

which matches the >9-day recovery timeframe empirically observed by Chang et al. [9]. Because the degradation rate is sampled from [0.01, 0.5] a.u.$^{-1}$ rather than fixed at one value, corresponding to real half-lives of approximately 12.5 minutes to 10.4 hours, the predicted heterogeneity-recovery time spans a wide range as well, from approximately 9.4 days at the stability threshold up to approximately 12.5 days across the full 2000 a.u. simulation window (2000 × 9 = 18,000 min). This range brackets the >9-day recovery timeframe empirically observed by Chang et al. [9], capturing both fast-switching networks that stabilize within hours and slow-switching networks whose recovery times match the biological benchmark.

### Regression and predictive models

*General framework.* All models take the circuit as the unit of analysis rather than the parameter set. The 1,000 parameter sets simulated per circuit are replicates within that circuit, so treating them as independent observations would understate standard errors by roughly the square root of the replicate count; every model therefore either aggregates to one value per circuit before fitting, or carries a variance estimator that respects the circuit grouping. Unless stated otherwise, the estimation sample is the 3,305 non-disjoint, non-null circuits carrying at least one feedback loop, drawn from the 3,411 non-isomorphic signed three-node motifs. Two design principles recur. First, because feedback categories differ systematically in density (median 5 edges for the pure categories versus 7 for the mixed) and density independently raises nearly every outcome studied here, each association is estimated both crude and density-adjusted, and both are reported so that confounding is visible rather than assumed away. Second, interpretation rests on effect sizes and interval estimates rather than on p-values, since a sample of several thousand circuits renders negligible differences significant.

*Topology features.* Predictors are drawn from a single feature table computed directly from the signed adjacency matrix, whose entries take values in {−1 (repress), 0 (absent), +1 (activate)} with rows as source and columns as target. The full set comprises 63 variables in five families: structure (18; the three self-regulation signs, the six directed edge signs, and summary counts of activating and repressing edges), degree (25; in-, out- and total degrees per gene together with their signed variants and distributional summaries), spectral (8; eigenvalue-based descriptors of the signed adjacency matrix), feedback taxonomy (6; counts of positive and negative loops of length one, two and three, a loop's sign being the product of its edge signs), and connectivity (6; strongly connected component profile and related indicators). Density (n_edges, the number of non-zero adjacency entries, self-loops included) serves throughout as the adjustment variable. Deprecated and redundant columns — proportional re-expressions such as edge_density, the binary has_* indicators, and the aggregate feedback totals — are excluded before fitting so that no predictor is an exact function of others in the same model. Three nested reduced sets are used in the parsimony comparison: density alone (1 feature), the feedback taxonomy alone (6 features), and their union (7 features). Every feature is named, grouped and defined in Supplementary Table S2, which is generated from the same function that builds the model input, so the table cannot drift from what the models actually receive.

**Supplementary Table S2.** The 63 topology features used in the regression and predictive models.

| Feature | Group | Description |
|---|---|---|
| $\mathrm{self\_reg}_1$ | Structure | Self-regulation sign of gene $x_1$ (diagonal; −1 repress / 0 none / +1 activate) |
| $\mathrm{self\_reg}_2$ | Structure | Self-regulation sign of gene $x_2$ (diagonal; −1 repress / 0 none / +1 activate) |
| $\mathrm{self\_reg}_3$ | Structure | Self-regulation sign of gene $x_3$ (diagonal; −1 repress / 0 none / +1 activate) |
| $\mathrm{edge}_{12}$ | Structure | Directed edge sign $x_1 \rightarrow x_2$ (−1 / 0 / +1) |
| $\mathrm{edge}_{13}$ | Structure | Directed edge sign $x_1 \rightarrow x_3$ (−1 / 0 / +1) |
| $\mathrm{edge}_{21}$ | Structure | Directed edge sign $x_2 \rightarrow x_1$ (−1 / 0 / +1) |
| $\mathrm{edge}_{23}$ | Structure | Directed edge sign $x_2 \rightarrow x_3$ (−1 / 0 / +1) |
| $\mathrm{edge}_{31}$ | Structure | Directed edge sign $x_3 \rightarrow x_1$ (−1 / 0 / +1) |

| Feature | Group | Description |
|---|---|---|
| $edge_{32}$ | Structure | Directed edge sign $x_3 \rightarrow x_2$ (−1 / 0 / +1) |
| n_activating | Structure | Count of activating entries (+1) in the matrix (incl. self-loops) |
| n_repressing | Structure | Count of repressing entries (−1) in the matrix (incl. self-loops) |
| n_edges | Structure | Total nonzero entries = number of edges incl. self-loops (network density) |
| n_self_activating | Structure | Number of activating self-loops (diagonal = +1) |
| n_self_repressing | Structure | Number of repressing self-loops (diagonal = −1) |
| n_self_none | Structure | Number of genes without self-regulation (diagonal = 0) |
| n_off_activating | Structure | Number of activating off-diagonal edges |
| n_off_repressing | Structure | Number of repressing off-diagonal edges |
| n_off_zero | Structure | Number of absent off-diagonal edges |
| $in\_deg\_pos_1$ | Degree | Activating incoming edges to $x_1$ (self-regulation excluded) |
| $in\_deg\_neg_1$ | Degree | Repressing incoming edges to $x_1$ |
| $out\_deg\_pos_1$ | Degree | Activating outgoing edges from $x_1$ |
| $out\_deg\_neg_1$ | Degree | Repressing outgoing edges from $x_1$ |
| $in\_deg\_total_1$ | Degree | Total incoming edges to $x_1$ (nonzero, self excluded) |
| $out\_deg\_total_1$ | Degree | Total outgoing edges from $x_1$ (nonzero, self excluded) |
| $net\_in\_deg_1$ | Degree | Net activating input to $x_1$ (in_deg_pos − in_deg_neg) |
| $in\_deg\_pos_2$ | Degree | Activating incoming edges to $x_2$ (self-regulation excluded) |
| $in\_deg\_neg_2$ | Degree | Repressing incoming edges to $x_2$ |
| $out\_deg\_pos_2$ | Degree | Activating outgoing edges from $x_2$ |
| $out\_deg\_neg_2$ | Degree | Repressing outgoing edges from $x_2$ |
| $in\_deg\_total_2$ | Degree | Total incoming edges to $x_2$ (nonzero, self excluded) |
| $out\_deg\_total_2$ | Degree | Total outgoing edges from $x_2$ (nonzero, self excluded) |
| $net\_in\_deg_2$ | Degree | Net activating input to $x_2$ (in_deg_pos − in_deg_neg) |
| $in\_deg\_pos_3$ | Degree | Activating incoming edges to $x_3$ (self-regulation excluded) |
| $in\_deg\_neg_3$ | Degree | Repressing incoming edges to $x_3$ |
| $out\_deg\_pos_3$ | Degree | Activating outgoing edges from $x_3$ |
| $out\_deg\_neg_3$ | Degree | Repressing outgoing edges from $x_3$ |
| $in\_deg\_total_3$ | Degree | Total incoming edges to $x_3$ (nonzero, self excluded) |
| $out\_deg\_total_3$ | Degree | Total outgoing edges from $x_3$ (nonzero, self excluded) |
| $net\_in\_deg_3$ | Degree | Net activating input to $x_3$ (in_deg_pos − in_deg_neg) |

| Feature | Group | Description |
|---|---|---|
| max_in_degree | Degree | Maximum in-degree across the three genes |
| max_out_degree | Degree | Maximum out-degree across the three genes |
| mean_in_degree | Degree | Mean in-degree across the three genes |
| mean_out_degree | Degree | Mean out-degree across the three genes |
| fb_pos_1node | Feedback | Positive 1-node feedback loops (activating self-loops) |
| fb_neg_1node | Feedback | Negative 1-node feedback loops (repressing self-loops) |
| fb_pos_2node | Feedback | Positive 2-node feedback loops (mutual pair, edge-sign product +1) |
| fb_neg_2node | Feedback | Negative 2-node feedback loops (mutual pair, edge-sign product −1) |
| fb_pos_3node | Feedback | Positive 3-node feedback loops (3-cycle, edge-sign product +1) |
| fb_neg_3node | Feedback | Negative 3-node feedback loops (3-cycle, edge-sign product −1) |
| n_scc | Connectivity | Number of strongly connected components (directed graph) |
| largest_scc_size | Connectivity | Size of the largest SCC (1–3; 3 = fully strongly connected) |
| is_strongly_connected | Connectivity | 1 if the whole graph is a single SCC, else 0 |
| n_wcc | Connectivity | Number of weakly connected components (edge direction ignored) |
| is_weakly_connected | Connectivity | 1 if the graph is weakly connected, else 0 |
| reachability | Connectivity | Fraction of ordered gene pairs ($i \neq j$) with a directed path $i \to j$ |
| trace | Spectral | Trace of the adjacency matrix (= sum of self-regulations) |
| det | Spectral | Determinant of the adjacency matrix |
| spectral_radius | Spectral | Largest eigenvalue magnitude, $\max\lvert\lambda\rvert$ |
| max_eigenvalue_real | Spectral | Largest real part among the eigenvalues |
| min_eigenvalue_real | Spectral | Smallest real part among the eigenvalues |
| sum_eigenvalue_real | Spectral | Sum of eigenvalue real parts (equals the trace) |
| eigenvalue_spread | Spectral | max_eigenvalue_real − min_eigenvalue_real |
| frobenius_norm | Spectral | Frobenius norm of the matrix (overall interaction strength) |

*Linear models.* Effects of feedback loops on continuous per-circuit outcomes — the median settling time, the pre-settling traversal rate, and the robust coefficient of variation (interquartile range divided by median) of settling time — are estimated by ordinary least squares on the six elementary loop counts, with HC3 heteroskedasticity-robust standard errors. Each coefficient reads as the change in the outcome per additional loop of that type. Each outcome is fitted twice, crude and with n_edges added, and both sets of coefficients are plotted together. A separate screening model for highly robust circuits regresses a binary top-decile indicator on n_edges, $n\_edges^2$ and one standardized feature at a time by logistic regression, reporting the log odds ratio per standard deviation with Holm correction across the features screened. For the variability outcome a third specification adding the settling-time level was fitted as a sensitivity analysis and is reported only as such: settling-time level is itself downstream of topology, so conditioning on it estimates direct rather than total effects, and the robust coefficient of variation carries the median in its denominator, which makes part of any level coefficient mechanical.

*Binomial and quasi-binomial models.* Robustness, i.e. the number of HR-enabling parameter sets out of 1,000, is modeled by binomial generalized linear models with a logit link. Per-loop effects are reported as the log odds per additional loop. Because circuit-level overdispersion is severe, confidence intervals are quasi-binomial, with standard errors inflated by the square root of the estimated dispersion parameter, which is reported alongside the coefficients. Category effects are summarized as density-adjusted rates by g-computation: the model $\text{logit}(p) \sim n\_edges + n\_edges^2 + \text{category}$ is fitted once, the entire sample is then counterfactually assigned to each category in turn, and the predicted probabilities are averaged over the observed density distribution, yielding a rate for each category at the common density profile. Confidence intervals and mirror-contrast p-values come from a network-level bootstrap that resamples circuits with replacement (500 replicates, percentile method); the model-based binomial standard error is not used for inference, as it ignores the between-circuit variance component and is far too small.

*Stratified density control.* Where the exposure and the outcome are both binary, density is controlled non-parametrically as well, by forming the two-by-two table within each stratum of n_edges and pooling by the Mantel–Haenszel estimator, with the Cochran–Mantel–Haenszel test for the pooled null and a Breslow–Day test of odds-ratio homogeneity across strata. Strata with a zero margin carry no information and are dropped; strata with a zero interior cell receive a Haldane–Anscombe correction of 0.5, which keeps the pooled estimate finite at the cost of shrinkage toward the null, so a corrected estimate is reported as a lower bound rather than as a point estimate. Rank associations are density-controlled by partial Spearman correlation on n_edges where a correlation rather than an odds ratio is the natural summary.

*Random-forest and gradient-boosting models.* Predictive ceilings and the parsimony comparison use tree ensembles. The settling-time ceiling compares a random-forest regressor (400 trees) and a gradient-boosting regressor (400 trees, learning rate 0.05, maximum depth 3, subsample fraction 0.8) against a mean-prediction baseline, all on the full topology feature set. The parsimony analysis fits a random-forest regressor (300 trees) to HR robustness and a random-forest classifier (300 trees) to HR existence, over the four nested predictor sets described above, plus a shuffled-target null fitted with 200 trees on the density-plus-feedback set. All remaining hyperparameters are scikit-learn defaults and were not tuned: unrestricted tree depth, minimum samples to split 2, minimum samples per leaf 1, bootstrap resampling enabled, and max_features of 1.0 for regressors or the square root of the predictor count for classifiers. Leaving these at their defaults is deliberate, since the analysis asks how much signal the features carry rather than how well a tuned model can be made to perform.

No single train–test split is used. All reported metrics are computed from pooled out-of-fold predictions under five-fold cross-validation, stratified on the class label for classification tasks and unstratified for regression. For the parsimony models the cross-validation is repeated 15 times with a freshly shuffled partition per repeat, the partition seed set to the repeat index and the estimator seed fixed at 42; the metric is computed once per repeat on that repeat's pooled out-of-fold predictions, and the mean and standard deviation across the 15 repeats are reported. The single-split cross-validation used for the predictive ceiling is seeded identically. Continuous targets are scored by the coefficient of determination and by the Spearman correlation between observed and out-of-fold predicted values; the existence classifier is scored by the area under the receiver operating characteristic curve and by the minority-class precision–recall area (non-HR circuits, prevalence approximately 0.10), the latter because the roughly 90/10 class balance makes the ROC area optimistic. Every model is benchmarked against the shuffled-target null fitted under the identical procedure. Variable importance, where reported, is permutation importance with 8 repeats computed on the fitted model, scored on the same metric as the corresponding cross-validation.

### Functional enrichment analysis with validated gene regulatory networks

The goal of this section is to identify genes that are involved in feedback loops in an experimentally validated gene regulatory network (GRN) and to quantify their functional enrichment, particularly with function annotations related to cell fate. The main components include the construction of a pooled network, the counting scheme for feedback loops, the development of density controlled null network model, and the function annotation analysis with cell-fate-related gene sets.

#### Construction of a pooled, signed, directed gene-regulatory network

##### Overview

All downstream analyses are built on a single GRN in which each gene is a node and each edge (arrow) records that one gene's product regulates another. Every edge carries two pieces of information. The first is its direction—it points from the regulator to the gene it controls (regulator → target). The second, where the evidence allows, is its sign—whether the regulator turns its target on (activation) or off (repression). Both components are required for identification of feedback loops, whose signs (positive or negative) depend on the signs of the edges in the loops.

To construct a comprehensive GRN for exploration of feedback loops, we sought to maximize two properties: broad coverage, so that as many real regulatory edges as possible are captured, and high signed coverage, so that as many edges as possible carry a known activation/repression sign. No single curated database delivers both. A transcription-factor database alone includes edges that correspond to transcriptional regulations and, by design, leaves out regulation that may act through signaling proteins in a pathway. We therefore combined three complementary, expert-curated databases, each providing directed and at least partly signed interactions, and combined them into a single, consistent set of human gene names: TRRUST [10] (literature-curated transcription-factor regulation), SIGNOR [11] (curated signaling interactions), and RegNetwork [12] (regulatory directions derived from the KEGG pathway database). We will describe the properties of each network data file and the assembled network in the next section.

##### Data sources

All three regulatory network data files for both human and mouse were downloaded and analyzed.

TRRUST v2 (Transcriptional Regulatory Relationships Unraveled by Sentence-based Text mining) is a database of transcription-factor → target gene regulation, hand-curated from published literature. Each interaction is labeled as activation, repression, or unknown (see a later section for the treatment of unknown interactions), and this label was used directly as the edge's sign. The human and mouse tables were obtained from the TRRUST website (grnpedia.org/trrust).

SIGNOR (Signaling Network Open Resource) is a database of expert-curated causal signaling events—one protein acting on another. We kept only protein-to-protein entries and discarded purely physical "form complex" (binding) records, which carry no information about activation or repression. The direction was taken as the curated source → target orientation, and the recorded mechanism was converted to a sign: mechanisms described as "up-regulates" became activation, "down-regulates" became repression, and anything else (for example cleavage or relocalization, where the on/off consequence is not specified) was treated as unknown. Human and mouse interactions were downloaded from the SIGNOR data service (signor.uniroma2.it).

RegNetwork provides regulatory directions derived from the KEGG pathway maps. From its direction tables, an activation symbol was scored as activation, an inhibition symbol as repression, and any other relation as unknown. Human and mouse files were taken from the RegNetwork download archive (regnetworkweb.org). Each source was loaded into one common table (regulator, target, sign, species, source), so that the pooling and sign-reconciliation steps that follow do not depend on which database an edge came from and new sources can be added later. Internally, signs were stored as +1 (activation), −1 (repression), and 0 (unknown, or—after pooling—conflicting) (see a later section for the treatment of conflicting interactions).

#### Pooling human and mouse gene regulations

To merge human and mouse interactions, each mouse gene was matched to its human ortholog using the Mouse Genome Informatics (MGI) mouse/human homology report [13]. We kept only high-confidence one-to-one matches, where exactly one mouse gene corresponds to exactly one human gene. Matches that were one-to-many or many-to-many (reflecting gene families with several similar copies, where the true counterpart is ambiguous) were discarded entirely, so that the merge could not invent regulatory edges—and therefore spurious feedback loops—between genes that are not truly equivalent. Of 21,929 homology groups, 18,784 gave clean one-to-one matches; the remaining 3,145 ambiguous groups were skipped.

All gene names were then standardized to human symbols (human names written in uppercase; mouse names converted through the one-to-one ortholog map, allowing for differences in capitalization between databases). Records whose regulator or target could not be resolved to a valid human gene symbol, including placeholder or non-gene tokens such as blanks, "NA", or "NULL", were dropped. Of 75,802 raw interaction records pooled across the three databases and both species, 72,254 were successfully placed on the human naming system, and 3,548 were discarded as unmatched or non-gene entries.

#### Assigning signs of edges

The same regulatory edge is often reported by more than one database or in both species. All records for a given regulator → target edge were therefore combined into a single merged edge, and that edge was given one agreed (consensus) sign from the evidence behind it: activation if every signed source agreed on activation, repression if every signed source agreed on repression, conflict if different sources disagreed (one says activation, another repression), and unknown if no source assigned a sign. Conflicting and unknown edges keep their direction but are treated as having no defined sign.

Each merged edge also retains a record of where it came from: which species and databases supported it, how many human and mouse records contributed, whether it was found in both human and mouse (a conserved edge), and how many independent databases reported it.

Two versions of the merged network were produced. The full network keeps every directed edge, including those whose sign is unknown or conflicting; it is used wherever only direction matters, such as categorizing feedback loops by size (one-, two-, and three-gene loops). The signed network keeps only edges with a single agreed sign (activation or repression), and it is used wherever every edge in a loop must have a definite sign. A loop's existence depends only on direction and is always well defined, but deciding whether a loop is positive or negative requires a sign on every edge, so that judgment is made only on the fully signed network.

##### Basic statistics of the assembled network

The full network contains 33,995 edges among 7,553 genes; 4,298 of these genes act as regulators (they have at least one outgoing edge) and 3,255 are regulated only (targets with no outgoing edge). A single agreed sign is available for 82.7% of edges (28,109 of them): 57.0% activation (19,379), 25.7% repression (8,730), 15.5% unknown (5,256), and 1.9% conflicting (630). Cross-species support is substantial: 19,982 edges are conserved (present in both human and mouse) and 2,152 are reported by more than one database. By source, the network draws 13,087 edges from TRRUST, 19,218 from SIGNOR, and 4,010 from RegNetwork (an edge can belong to more than one source).

Using the TRRUST database alone gives signed coverage of only about 58% and a small loop-containing core; adding the signaling and pathway databases raises signed coverage to 82.7% and enlarges the loop-containing core roughly five-fold (from about 517 to 2,521 genes). This shows that using the combined sources had a significant impact on the comprehensiveness of the derived network. The three-database union is the default network used throughout.

The network also contains the raw material needed for feedback analysis. There are 311 self-loops (genes that regulate themselves—one-gene feedback; among the signed ones, 223 are self-activating and 47 self-repressing) and 758 mutually regulating gene pairs (two genes that each regulate the other—candidate two-gene loops). To find the part of the network capable of feedback at all, we computed its strongly connected components (SCCs): groups of genes so wired that, starting from any gene in the group and following edges in their direction, one can reach every other gene in the group and return. Together, 2,538 genes lie in a multi-gene feedback component, and 311 carry self-loops. Their union is a set of 2,588 genes. As an example of known feedback regulations, the Notch-pathway two-gene negative loop is recovered intact and conserved across species—HES1 represses RBPJ while RBPJ activates HES1—as are the repression of TP53 by MDM2 and the activation of NANOG by POU5F1 and by itself, both central to stem-cell pluripotency.

#### Counting feedback loops and building a per-gene feedback profile

##### Overview

This step turns the network into the single quantity the function analyses depend on: for each gene, how many feedback loops it takes part in, broken down by loop size and by whether each loop is positive or negative. This involves three operations on the full network from the previous subsection: narrowing to the genes that can lie on a loop, listing all the short loops exhaustively, and labeling each loop. Next, the per-gene counts are tallied. We refer to a gene's set of counts as its feedback profile.

##### Narrowing to the genes that can lie on a loop

Only a gene that sits on a directed loop can take part in feedback, so the search was confined to the feedback-capable part of the network rather than all 7,553 genes. This part is the union of the genes in a multi-gene strongly connected component (the feedback regions defined above) and the genes with self-loops, and it comprises 2,588 genes. Restricting the search to these genes makes an exhaustive loop count feasible without missing anything, because no loop can pass through a gene outside this set.

##### Listing every short loop

Every simple directed loop of one to four genes was listed (simple means that the loop visits each gene only once and so represents a single distinct feedback circuit). We used the loop-finding routine in the networkx library [14], which returned 51,411 such loops. The size was capped at four genes for two reasons: the number of longer loops explodes combinatorially and would be intractable, whereas loops of one to four genes already span the biologically meaningful cases—a gene acting on itself (one gene), a mutually regulating pair (two genes), and the short three-gene circuits—while still allowing four-gene loops as an outer limit. Because each loop is simple, a gene is counted at most once per loop.

##### Labeling each loop by size and by sign

Each loop was described on two axes. Its size is simply the number of genes it contains (one to four). Its sign captures whether the loop, taken as a whole, is positive or negative, and is obtained by multiplying together the activation/repression signs of its edges (each activation counts as +1 and each repression as −1). A loop is positive if the product is +1 and negative if the product is −1. If any edge in the loop has an unknown or conflicting sign, the product is undefined and the loop is recorded as unsigned: it still counts toward size but not toward sign, matching the unknown-sign policy above.

It is worth noting that multiplying the signs means that a double-negative loop (two genes that repress each other) yields net *positive* and behaves as a bistable switch (the system settles into one of two stable states). So "positive feedback" includes both mutual activation and mutual repression; a loop is negative only when it contains an odd number of repressions. Accordingly, two-gene loops were further split into mutual activation ("PP"), mutual repression ("NN", net positive), and mixed ("PN", net negative), and three-gene loops were split by their number of repressions (zero or two repressions give a positive loop; one or three give a negative loop).

Across all sizes, 10,705 loops are positive, 9,799 are negative, and 30,907 are unsigned (39.9% of loops are signed). The fraction with a known sign is high for the smallest loops and falls as loops grow: 86.8% of one-gene loops (223 positive, 47 negative, 41 unsigned; 311 total), 69.1% of two-gene loops (342 positive, 182 negative, 234 unsigned; 758 total), 51.5% of three-gene loops (931 positive, 907 negative, 1,732 unsigned; 3,570 total), and only 38.2% of four-gene loops (9,209 positive, 8,663 negative, 28,900 unsigned; 46,772 total). Two-gene loops comprise 247 mutual-activation, 95 mutual-repression, and 182 mixed; the signed three-gene loops split by number of repressions as 480 (none), 800 (one), 451 (two), and 107 (three). Positive and negative loops occur in similar numbers overall (10,705 versus 9,799).

#### The per-gene feedback profile

For each gene, its participation was tallied across all loops into a profile that records: the count in each size-and-sign category (for example, positive three-gene loops); the totals across categories (total loops, and the numbers that are positive, negative, unsigned, and signed); the size of the largest loop the gene belongs to; and basic connectivity measures. Connectivity is summarized by a gene's degree—the number of edges touching it, separated into incoming edges (regulators acting on the gene) and outgoing edges (genes it regulates), together with the size of its feedback component. Of 7,553 genes, 1,863 take part in at least one feedback loop, 1,420 in at least one positive loop, and 1,266 in at least one negative loop. These profiles, with their connectivity measures, are the per-gene input to the function-enrichment analyses.

In function enrichment analyses described in subsequent sections, genes' feedback profiles will be used to quantify the association with annotated gene functions. However, controlling for connectivity in such analyses is necessary for two reasons. First, because the fraction of signed loops falls with size, the cleanest sign information lives in the one- to three-gene loops, which is where the sign-based analyses are focused. Second, the raw loop count is dominated by a gene's connectivity: the most heavily connected genes (hubs) sit in enormously many loops simply by virtue of their many edges. TP53, for instance, takes part in 11,559 loops and has 507 edges, and other hubs are similarly loop-rich (NOTCH1: 1,154 loops, 139 edges; MDM2: 1,179 loops, 114 edges; HES1: 502 loops, 72 edges), whereas a well-known but less-connected regulator such as NANOG sits in only 76 loops (47 edges). Because loop count rises with connectivity, the number of loops a gene is in cannot be compared fairly between genes without accounting for how connected each gene is. This motivates the connectivity-matched (degree-preserving) null model introduced later.

#### Defining the biological functions and a preliminary, connectivity-aware enrichment test

##### Overview

The two previous subsections produce, for every gene, a measure of how much feedback it takes part in. This step asks the biological question: are genes that sit in feedback loops disproportionately involved in particular functions—specifically stem/progenitor-cell maintenance and cellular development? To do so it must first decide, from an external and unbiased source, which genes count as "stem/progenitor" genes and which count as "cellular-development" genes, and then test whether feedback-participating genes are over-represented among them. Because the previous subsection showed that the number of loops a gene is in rises steeply with its connectivity, and because well-connected hub genes also tend to be the most heavily annotated, this step introduces connectivity (degree) controls and treats its results as a preliminary test that motivates the stricter test that follows.

##### Defining the function gene sets from the Gene Ontology database

Gene sets were defined using the Gene Ontology (GO) [15,16], a standardized, expert-maintained vocabulary of biological functions in which terms are arranged from general to specific and linked by defined hierarchical relationships. For example, "stem-cell population maintenance" is a more specific case of broader developmental terms. We used the Biological Process branch. Each function of interest was defined by a small number of anchor GO terms and then expanded to include every more specific term beneath them (its descendants), so that lineage- or tissue-specific child terms are captured automatically rather than having to be listed by hand. Stem/progenitor was anchored on stem-cell population maintenance, somatic stem-cell maintenance, stem-cell division, and stem-cell proliferation. Cellular development was anchored on the single broad term "cellular developmental process"—the development of cells, a wide-ranging program covering the many specific ways in which cells acquire their specialized identities and states; because the anchor is expanded to all of its descendants, this set automatically subsumes numerous more specific developmental sub-processes, a nested structure we exploit later. As references, homeostasis was anchored on homeostatic process and tissue homeostasis, and, as a positive control, cell cycle on the cell-cycle term [17,18].

The expansion followed the true biological path through the hierarchy, using both kinds of GO relationship, i.e. "is a type of" (is_a) and "is part of" (part_of), rather than the more common shortcut of following only "is a type of" links. This matters because much of the developmental and cell-cycle hierarchy is connected through "is part of" relationships, so the shortcut silently undercounts the relevant genes; following the true path expands the cellular-development branch to thousands of descendant terms, far more than the shortcut would capture. GO term definitions and relationships were read with the goatools library [19], and gene-to-GO annotations were retrieved through the MyGene.info service [20].

#### Restricting to trustworthy annotations

Each gene–function annotation in GO carries an evidence code recording how it was established. The primary analysis used only curated annotations—those backed by experiments (for example, direct assay, or the effect of a mutation) or by a curator reading a paper—and excluded annotations assigned automatically by computational or evolutionary inference. Automatic annotations were excluded because they are heavily biased toward broadly studied housekeeping genes and would inflate apparent associations; a secondary analysis that includes all evidence codes was also computed as a sensitivity check.

One practical problem is that the curated GO stem-cell set is very small (approximately 28 genes), too few to give the test adequate statistical power. We therefore supplemented the stem/progenitor set with widely used, curated stem-cell gene signatures from the Molecular Signatures Database (MSigDB) [21,22] to construct broad embryonic-stem-cell-core and pluripotency-network panels. Target-gene sets of the master pluripotency regulators NANOG, OCT4, and SOX2 (the Wong, BenPorath, Bhattacharya, Ramalho, and Mueller signatures) were pooled, and this raised the stem/progenitor set to roughly 1,200 network genes. This enrichment applies only to the stem/progenitor category; cellular development and the reference categories are defined from GO alone.

#### The enrichment test and its connectivity controls

For each gene, the feedback feature tested was simply whether it takes part in at least one signed feedback loop of three or fewer genes, the size range consistent with our mathematical models. Enrichment of feedback-feature genes in each function was then estimated in three measures/estimates, two of which were basic treatments of hub confound, which will be replaced with a better approach described in the next section. The measure of association is the odds ratio, reported throughout as its natural logarithm (the log-odds ratio): a value of 0 means feedback genes are no more likely than other genes to have the function, positive values mean they are more likely, and larger values mean stronger enrichment (as a rough guide, +0.7 corresponds to roughly a doubling of the odds). The three estimates were: (i) a raw log-odds ratio with its Fisher exact-test p-value, applying no connectivity control; (ii) a Mantel–Haenszel log-odds ratio, which pools the association after splitting genes into five equal-sized bins by connectivity (degree quintiles) so that feedback and non-feedback genes are compared within similar-connectivity groups; and (iii) a logistic regression of function on the feedback feature together with the logarithm of connectivity, which controls for degree as a continuous variable. Each test was run on two gene sets: all GO-annotated genes, and the stricter set of annotated regulators (genes with at least one outgoing edge).

##### Associations from the naïve test

Among annotated regulators using curated evidence (1,036 of them carry the signed-feedback feature), the raw enrichment is large and highly significant for cellular development (log-odds +0.79 (19.5% of feedback genes are cellular-development genes versus 9.9% of the rest; Fisher p $\approx 2\times10^{-14}$)) and modest but significant for stem/progenitor (+0.28; 19.8% versus 15.7%; p $\approx 3\times10^{-3}$). Crucially, these raw associations shrink sharply once connectivity is controlled: the connectivity-matched (Mantel–Haenszel) estimates fall to +0.14 for cellular development (95% CI −0.10 to +0.39) and +0.10 for stem/progenitor (95% CI −0.13 to +0.32)—cellular development retaining a positive but attenuated residual and stem/progenitor's residual no longer distinguishable from zero. In other words, most of the apparent link between feedback and function is explained by the fact that hubs are both feedback-rich and heavily annotated. The two references behave as expected: homeostasis shows no positive association (connectivity-matched log-odds −0.33), and the cell-cycle control remains clearly enriched even after connectivity control (+0.49; 95% CI +0.17 to +0.81), which is biologically appropriate because the cell cycle is itself a feedback-controlled cellular system and therefore serves as a positive control that the method detects genuine feedback-related functions.

Two further descriptive analyses were run on the same regulator set. Splitting the feedback feature by loop size showed that, for cellular development, the connectivity-matched enrichment is carried by the larger short loops rather than by self-regulation (about −0.13, +0.20, and +0.23 for one-, two-, and three-gene loops). Splitting it by loop sign showed that positive and negative feedback give almost identical enrichments for every function (differences of at most about 0.06 in log-odds), a first piece of evidence supporting our conclusion that it is participation in feedback, not its sign, that tracks function.

Splitting genes into five connectivity bins is only a coarse control: genes within a bin still differ in connectivity, and binning cannot ask whether a particular gene sits in more feedback than its own specific wiring would predict. Answering that requires holding each gene's exact connectivity fixed while randomizing the rest of the network. i.e. the connectivity-preserving null model developed next, which converts these borderline preliminary estimates into a definitive test.

#### A connectivity-preserving null model

##### Overview

This step delivers the definitive verdict. Instead of grouping genes into connectivity bins, it holds each gene's exact wiring pattern fixed and asks a sharper question: does a gene sit in *more* signed feedback loops than would be expected from its connectivity alone? The comparison is made against many randomized networks that preserve every gene's connectivity but scramble everything else, so that any excess of feedback a gene shows above this random expectation cannot be an artifact of how many edges it happens to have.

##### Randomizing the wiring while preserving each gene's connectivity

A null model is a randomized version of the network used as a yardstick for what "no special structure" looks like. Here the randomization is a degree-preserving edge swap [23]: pairs of edges are repeatedly picked and their endpoints exchanged (edge A→B and C→D become A→D and C→B), which rewires the network while leaving every gene with exactly the same number of edges it started with. Two refinements make the yardstick strict. First, swaps are performed separately within each sign class (activating, repressing, and unknown edges are only ever exchanged with edges of the same sign), so that each gene keeps not just its total number of incoming and outgoing edges but its exact counts of activating and repressing edges of each direction. Second, self-loops are held fixed. The result is a network in which each gene's complete connectivity signature is identical to the real one, but the specific partners (and therefore the feedback-loop structure) are randomized. We generated 200 such randomized networks.

##### Measuring feedback beyond connectivity with a z-score

In the real network and in each randomized one, we recounted every gene's participation in signed two- and three-gene feedback loops (one-gene self-loops are unchanged by this randomization and so are excluded). For each gene this gives an observed count and a distribution of counts expected under the null, summarized as a z-score: how many standard deviations the observed count lies above (or below) the average of the randomized networks. A z-score of 0 means the gene sits in exactly as much feedback as its connectivity predicts; a large positive z-score means it sits in far more. Genes with a z-score of 2 or greater were labeled feedback-rich (493 such genes fall within the annotated-regulator set used for testing). Because "feedback-rich" is already defined relative to each gene's own connectivity, a simple comparison of feedback-rich versus the rest is connectivity-controlled by construction. Therefore, no further adjustment is needed. Separate z-scores were also computed for positive and for negative feedback.

##### Enrichment of cell-fate related functions

Testing enrichment of feedback-rich genes in each function (annotated regulators, curated evidence) gives the following connectivity-controlled log-odds ratios. Cellular development is enriched at +0.37 (95% CI +0.11 to +0.63; 16.4% of feedback-rich genes are cellular-development genes versus 12.0% of the rest; Fisher $p = 6.9\times10^{-3}$). Stem/progenitor is enriched at +0.27 (95% CI +0.03 to +0.50; 20.3% versus 16.3%; p = 0.033). Homeostasis shows no association (−0.12; p

= 0.66), and the cell-cycle control had the strongest association (+0.83; 95% CI +0.53 to +1.14; p = $3.7\times10^{-7}$), as expected for a process that is itself a feedback-controlled system.

##### Positive and negative feedback are indistinguishable

Using the separate positive- and negative-feedback z-scores, both kinds of feedback are individually enriched for cellular development and to essentially the same degree: the log-odds are +0.36 for positive and +0.35 for negative feedback (difference +0.01, p = 0.96). The same holds for stem/progenitor (+0.32 for positive and +0.29 for negative; difference +0.03, p = 0.89). In no case do positive and negative feedback differ significantly. Loop sign therefore does not partition function in our tests.

The cellular-development result is also robust to plausible variations of the analysis, including rebuilding the network from the transcription-factor database alone, restricting the feedback network to transcription factors only, and removing proliferation-associated genes; the enrichment persists in each case.

#### Enrichment of specific cell-differentiation programs

The previous subsection establishes that feedback-rich genes are enriched for cellular development. However, cellular development is a broad umbrella covering many specific processes, so we asked whether the feedback association is spread evenly across development or concentrated in a particular part of it. Because the cellular-development gene set was built by expanding a single anchor term to all of its GO descendants, we can reuse that same hierarchy, and the null-model z-scores already computed above, with no new randomization, to walk from broad developmental terms down to specific ones and see where the enrichment actually lives. This analysis refines an already-established association rather than testing a new confirmatory hypothesis.

Each GO term along the path was expanded to its own descendant-closed gene set (true path, is_a + part_of), and enrichment of feedback-rich genes ($z \geq 2$) was recomputed within the same universe of annotated regulators, reported as a connectivity-controlled log-odds ratio and split by feedback sign.

At the broad ancestor developmental process (942 genes), feedback-rich genes are *not* enriched at all (log-odds +0.05; 95% CI −0.16 to +0.27; p = 0.62). One level down, at cellular developmental process—the function confirmed above (472 genes)—the enrichment appears (+0.37; 95% CI +0.11 to +0.63; p = $6.9\times10^{-3}$). Descending further within it, the signal is fully accounted for by one specific descendant: cell differentiation, the process by which cells acquire specialized identities, is enriched at +0.37 (95% CI +0.11 to +0.63; $p = 6.9\times10^{-3}$)—numerically identical to its parent, because in this network the two terms are supported by the same set of regulators—whereas the neighboring branch cell development is not significant (+0.20; 95% CI −0.13 to +0.54; p = 0.24). In short, the developmental feedback signal hypothesized and confirmed as cellular development is, on closer inspection, specifically a cell-differentiation signal: absent from the broad developmental ancestor and concentrated precisely at cell differentiation and its immediate parent. That it is null one level higher rules out a generic "developmental gene" or hub artifact that would otherwise smear across the whole branch.

We made two interesting observations regarding the distributions of the enrichment. First, the cell-differentiation signal is not driven by any single cell lineage: the specific differentiation branches tested—neuron, leukocyte, myeloid, muscle, and hematopoietic-stem-cell differentiation—are each individually non-significant and underpowered (as few as 6 to 121 genes), indicating a broad cell-differentiation program rather than one cell type. Second, the enrichment sits in the differentiation processes themselves more than in their regulation: the "regulation of cell differentiation" sub-branch is weaker and non-significant (+0.21; p = 0.17). Within that sub-branch there is a non-significant hint that negative feedback loads onto negative regulation of differentiation (negative-feedback log-odds +0.60 versus positive-feedback +0.21), but this rests on few genes across many slices and is reported as hypothesis-generating only rather than a conclusion.

#### Curated cell-fate circuits and their biological contexts

The feedback statistics reported above are network-wide. To connect them to concrete biology, we assembled a curated catalog of well-characterized cell-fate circuits, each of which is contained in the pooled network, corresponds to a nameable progenitor or differentiation system, and has direct experimental support in the literature. For each circuit, the signed feedback loops lying entirely within its gene set were counted and summarized by the frustration index $n_{neg} / (n_{pos} + n_{neg})$, which assigns the circuit to one of five polarity groups: pure positive (0), mixed positive-dominant (< 0.5), balanced (0.5), mixed negative-dominant (> 0.5), and pure negative (1). Supplementary Table S3 lists the circuits, the biological system each represents, and the primary literature supporting that assignment. All eleven circuits fall into the first four groups; no curated fate circuit was purely negative. For six of the eleven circuits we also identified published single-cell evidence that the circuit's genes are expressed bi- or multimodally and that the modes correspond to interconverting subpopulations rather than fixed cell types; these are cited in the final column of Supplementary Table S3.

**Supplementary Table S3.** Curated cell-fate circuits, the progenitor or differentiation systems they represent, and their supporting literature.

| Feedback group | Circuit | Biological system | System evidence | Inter-conversion evidence |
|---|---|---|---|---|
| Pure positive | ICM vs trophectoderm | First blastocyst lineage split (epiblast/ICM vs trophectoderm) | [24] | [25] |
| Pure positive | Retinoic-acid receptors | RA-driven neural differentiation; PPARG-driven adipogenic differentiation | [26,27] | — |
| Pure positive | EMT switch | Neural-crest delamination / gastrulation EMT | [28] | [29,30] |
| Mixed, positive-dominant | Pluripotency core | Embryonic stem cell / iPSC self-renewal | [31] | [32,33] |
| Mixed, positive-dominant | Hematopoietic GATA1–PU.1 | Myeloid progenitor lineage choice | [34] | [9] |
| Balanced | Neural progenitor Notch-HES | Neural progenitor maintenance / lateral inhibition | [35] | — |
| Mixed, negative-dominant | Wnt/β-catenin–AXIN2 | Intestinal crypt stem cell self-renewal | [36] | — |
| Mixed, negative-dominant | TGF-β/BMP–SMAD | Osteogenic differentiation of mesenchymal/myogenic progenitors | [37] | — |
| Mixed, negative-dominant | Notch-HES-HEY (broad) | Notch lateral inhibition across progenitor compartments | [38] | [39] |
| Mixed, negative-dominant | p53–MDM2 | Stem/progenitor quiescence and reprogramming barrier | [40] | — |
| Mixed, negative-dominant | Myogenic (MyoD) | Muscle satellite-cell specification and self-renewal | [41] | [42,43,44] |

### Supplementary Figures

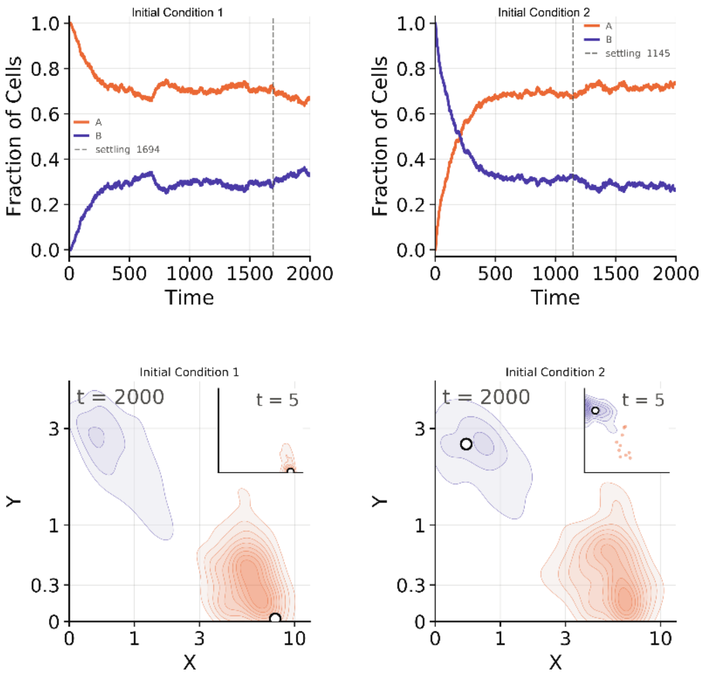

**Figure S1. An example of heterogeneity restoration**. A representative parameter set for a gene network containing a mutual inhibition circuit (toggle switch) was used as an example. Two initial conditions were chosen (left and right), one from deterministic ODE simulation, and the other from the mode of the first SODE simulation run that was further from the initial conditions. Top plots show dynamics of fractions of two cell types defined by a two-population Gaussian Mixture Model (GMM). Vertical lines indicate settling times. Bottom plots show the distributions of cells in gene expression space at two representative timepoints.

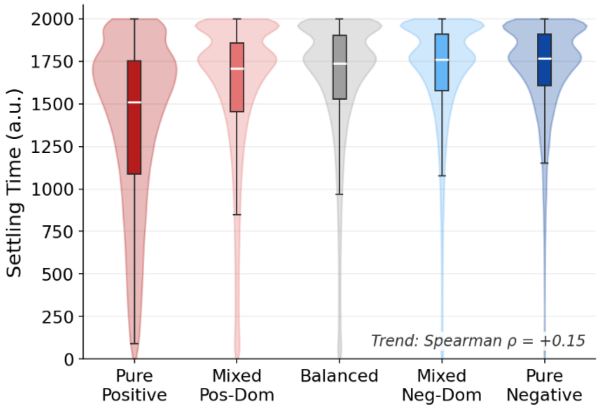

**Figure S2. Distributions of parameter-wise median settling time for 5 categories of networks**. Networks were categorized based on the number of positive and negative feedback loops. Basal level of noise was used for all simulations. The settling time was defined as the last time point at which this fraction differed from its steady-state value (the mean value of its fraction over the final 200 time units of the simulation) by more than 5% in absolute cell fraction.

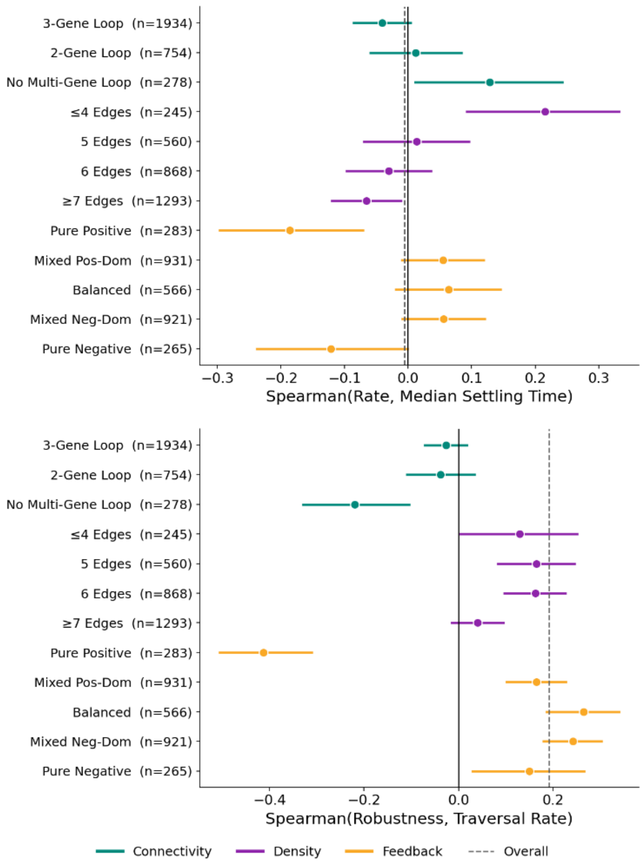

**Figure S3. Correlations of pre-settling traversal rate, settling time, and robustness**. Rows show subgroups of 2966 networks that were independently created. Spearman correlation coefficients were calculated based on network-wise median traversal rate and either settling time (top) or HR robustness (bottom). The horizontal bars are 95% CIs from the Fisher z-transform $z = \text{atanh}(\rho)$, $SE = 1/\sqrt{(n - 3)}$, then ±1.96 SE transformed back. The dashed line is the overall correlation across all networks.

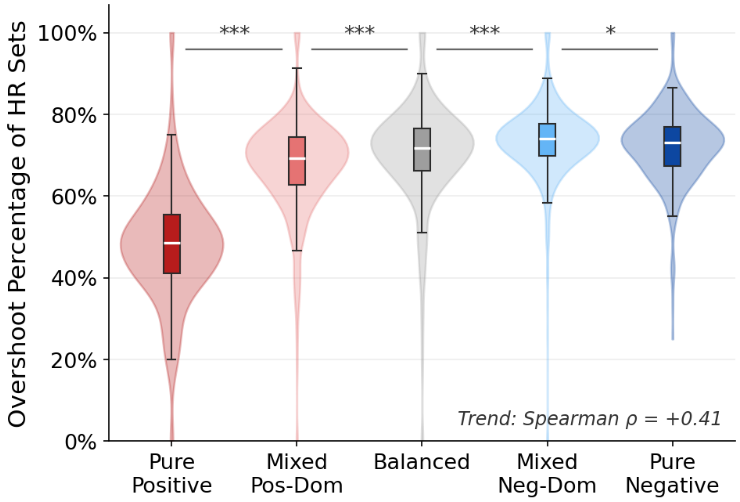

**Figure S4. Distributions of percentages of parameter sets with overshoot dynamics for 5 categories of networks**. Networks were categorized based on the number of positive and negative feedback loops. Basal level of noise was used for all simulations. An overshoot behavior was defined as the case when a fraction of the non-initial cell type reached a peak value at least 10% higher than its final value. *: $p < 0.05$. ***: $p < 0.001$.

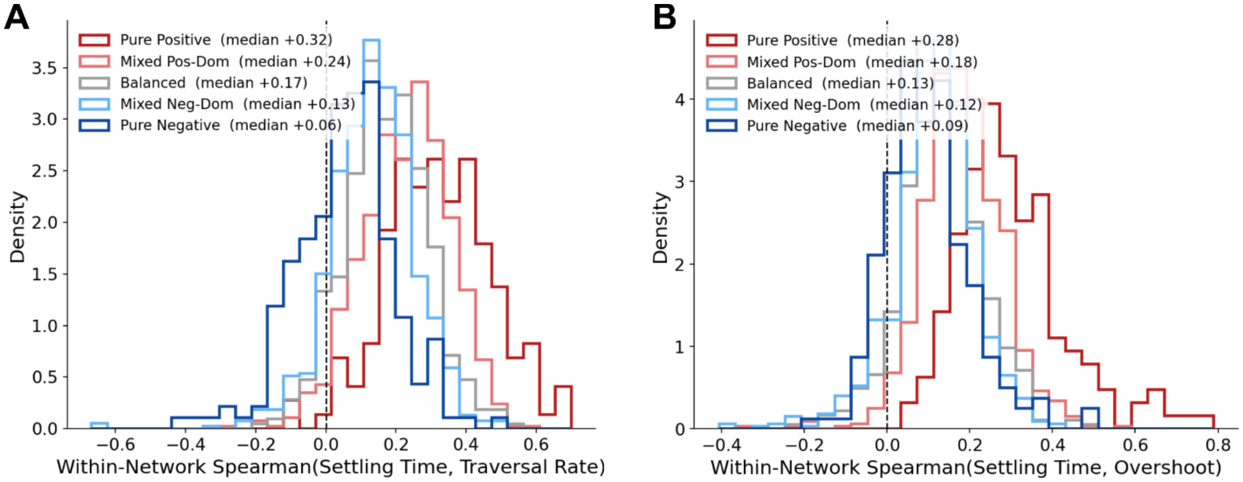

**Figure S5. Distributions of within-network correlations**. Parameter-set-wise settling time, traversal rate, and overshoot dynamics were used to compute within-network Spearman correlation coefficients. **A**. Correlation between settling time and traversal rate. **B**. Correlation between settling time and overshoot dynamics (similar to a scaled result from the Mann–Whitney U test).

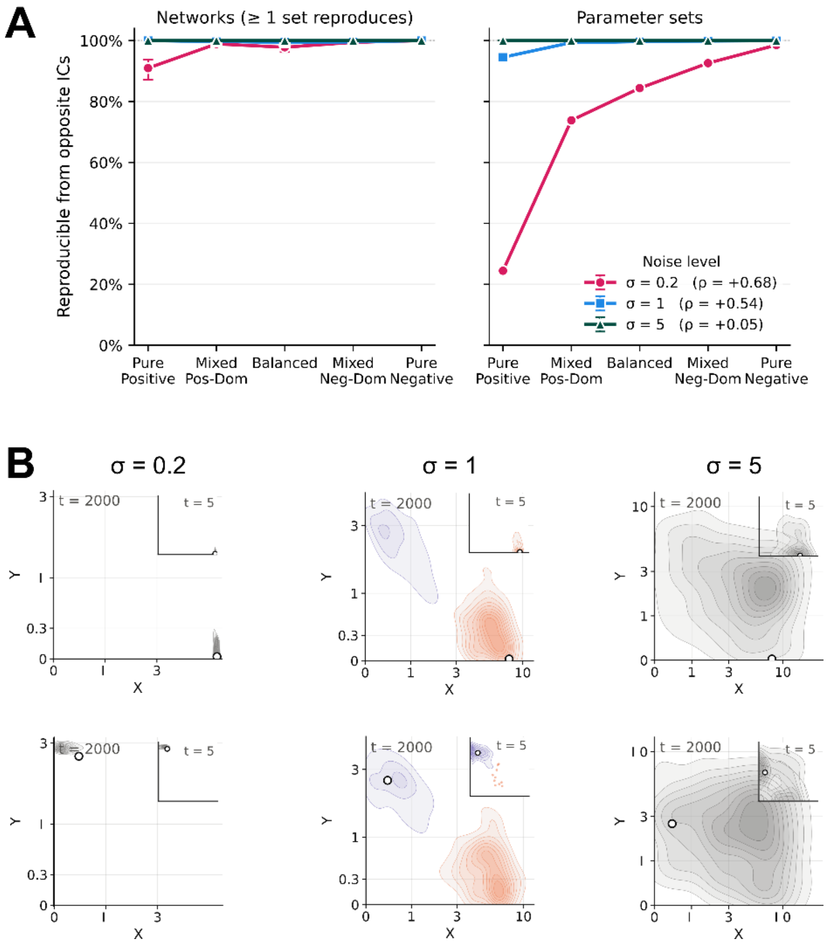

**Figure S6. Perturbation of heterogeneity restoration induced by changes of noise level**. **A**. Reproducibility of multimodal gene expression patterns in cell populations at different noise levels and feedback categories when two initial conditions (ICs) were used. Left plot shows network-wise reproducibility. Right plot shows parameter-set-wise reproducibility. Only parameter sets that passed filters before the opposite initial condition tests were included. **B**. Illustration of two scenarios of HR failures due to change in noise level. A representative parameter set for a gene network containing a mutual inhibition circuit (toggle switch) was used as an example. At the basal noise level (middle), the parameter set was HR-enabling. At the low and high noise levels (left and right), neither initial condition produces a multimodal pattern. Contour maps show distributions of cells in gene expression space with types defined by a two-population Gaussian Mixture Model (GMM) at two representative timepoints.

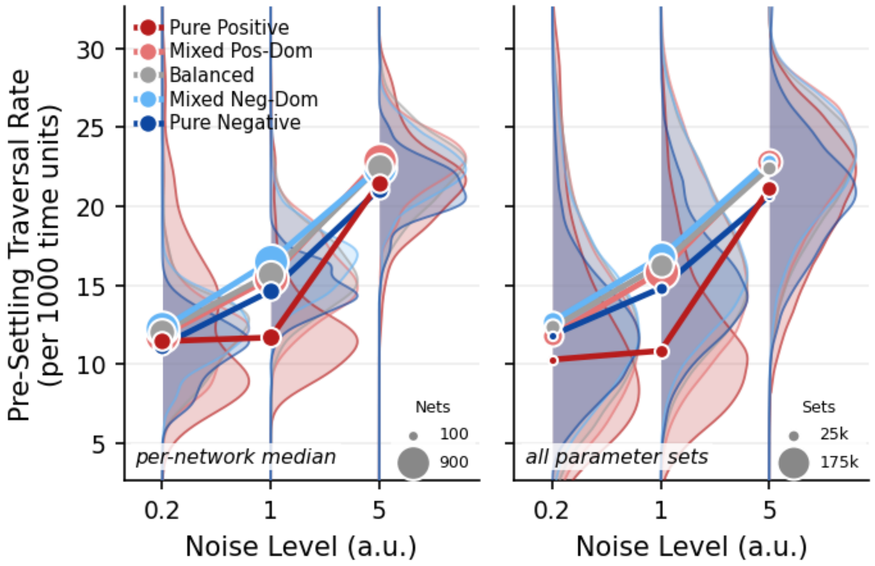

**Figure S7**. **Distributions of pre-settling traversal rate at three noise levels**. Left: Distributions of median pre-settling traversal rate for indicated categories of networks. KDE plots show distributions of network medians. Dots show means of median pre-settling traversal rates for each category. Right: Distributions of parameter-set-wise pre-settling traversal rate for indicated categories of networks. KDE plots show distributions of pre-settling traversal rates of all parameter sets in each category. Dots show medians of pre-settling traversal rates for all parameter sets in each category.

### References

1. Limpert E, Stahel WA, Abbt M. Log-normal distributions across the sciences: keys and clues. *BioScience*. 2001;51(5):341. doi:10.1641/0006-3568(2001)051[0341:LNDATS]2.0.CO;2.

2. Devroye L. Non-Uniform Random Variate Generation. New York: Springer-Verlag; 1986.

3. Holder J, Mohammed S, Barr FA. Ordered dephosphorylation initiated by the selective proteolysis of cyclin B drives mitotic exit. *eLife*. 2020;9:e59885. doi:10.7554/eLife.59885.

4. Schwanhäusser B, Busse D, Li N, Dittmar G, Schuchhardt J, Wolf J, Chen W, Selbach M. Global quantification of mammalian gene expression control. *Nature*. 2011;473(7347):337–342. doi:10.1038/nature10098.

5. Elowitz MB, Leibler S. A synthetic oscillatory network of transcriptional regulators. *Nature*. 2000;403(6767):335–338. doi:10.1038/35002125.

6. Gardner TS, Cantor CR, Collins JJ. Construction of a genetic toggle switch in *Escherichia coli*. *Nature*. 2000;403(6767):339–342. doi:10.1038/35002131.

7. Nash P, Tang X, Orlicky S, Chen Q, Gertler FB, Mendenhall MD, Sicheri F, Pawson T, Tyers M. Multisite phosphorylation of a CDK inhibitor sets a threshold for the onset of DNA replication. *Nature*. 2001;414(6863):514–521. doi:10.1038/35107009.

8. Goldbeter A, Koshland DE. An amplified sensitivity arising from covalent modification in biological systems. *Proceedings of the National Academy of Sciences of the United States of America*. 1981;78(11):6840–6844. doi:10.1073/pnas.78.11.6840.

9. Chang HH, Hemberg M, Barahona M, Ingber DE, Huang S. Transcriptome-wide noise controls lineage choice in mammalian progenitor cells. *Nature*. 2008;453(7194):544–547. doi:10.1038/nature06965.

10. Han H, Cho JW, Lee S, Yun A, Kim H, Bae D, Yang S, Kim CY, Lee M, Kim E, Lee S, Kang B, Jeong D, Kim Y, Jeon HN, Jung H, Nam S, Chung M, Kim JH, Lee I. TRRUST v2: an expanded reference database of human and mouse transcriptional regulatory interactions. *Nucleic Acids Research*. 2018;46(D1):D380–D386. doi:10.1093/nar/gkx1013.

11. Lo Surdo P, Iannuccelli M, Karis K, Meo E, Omidi P, Tosoni M, Graziosi S, Panni S, Fiorillo MT, Licata L, Sacco F, Gyori BM, Perfetto L. SIGNOR 4.0: the 2025 update with focus on phosphorylation data. *Nucleic Acids Research*. 2025;gkaf1237. doi:10.1093/nar/gkaf1237.

12. Liu ZP, Wu C, Miao H, Wu H. RegNetwork: an integrated database of transcriptional and post-transcriptional regulatory networks in human and mouse. *Database (Oxford)*. 2015;2015:bav095. doi:10.1093/database/bav095.

13. Baldarelli RM, Smith CL, Ringwald M, Richardson JE, Bult CJ; Mouse Genome Informatics Group. Mouse Genome Informatics: an integrated knowledgebase system for the laboratory mouse. *Genetics*. 2024;227(1):iyae031. doi:10.1093/genetics/iyae031.

14. Hagberg AA, Schult DA, Swart PJ. Exploring network structure, dynamics, and function using NetworkX. In: *Proceedings of the 7th Python in Science Conference (SciPy 2008)*. Pasadena, CA; 2008. p. 11–15.

15. Ashburner M, Ball CA, Blake JA, Botstein D, Butler H, Cherry JM, Davis AP, Dolinski K, Dwight SS, Eppig JT, Harris MA, Hill DP, Issel-Tarver L, Kasarskis A, Lewis S, Matese JC, Richardson JE, Ringwald M, Rubin GM, Sherlock G. Gene ontology: tool for the unification of biology. The Gene Ontology Consortium. *Nature Genetics*. 2000;25(1):25–29. doi:10.1038/75556.

16. The Gene Ontology Consortium; Aleksander SA, Balhoff J, Carbon S, et al. The Gene Ontology knowledgebase in 2023. *Genetics*. 2023;224(1):iyad031. doi:10.1093/genetics/iyad031.

17. Kapuy O, He E, López-Avilés S, Uhlmann F, Tyson JJ, Novák B. System-level feedbacks control cell cycle progression. *FEBS Letters*. 2009;583(24):3992–3998. doi:10.1016/j.febslet.2009.08.023.

18. Dragoi C, Tyson JJ, Novák B. Newton's cradle: cell cycle regulation by two mutually inhibitory oscillators. *Mathematical Biosciences*. 2024;377:109291. doi:10.1016/j.mbs.2024.109291.

19. Klopfenstein DV, Zhang L, Pedersen BS, Ramírez F, Warwick Vesztrocy A, Naldi A, Mungall CJ, Yunes JM, Botvinnik O, Weigel M, Dampier W, Dessimoz C, Flick P, Tang H. GOATOOLS: a Python library for Gene Ontology analyses. *Scientific Reports*. 2018;8:10872. doi:10.1038/s41598-018-28948-z.

20. Xin J, Mark A, Afrasiabi C, Tsueng G, Juchler M, Gopal N, Stupp GS, Putman TE, Ainscough BJ, Griffith OL, Torkamani A, Whetzel PL, Mungall CJ, Mooney SD, Su AI, Wu C. High-performance web services for querying gene and variant annotation. *Genome Biology*. 2016;17:91. doi:10.1186/s13059-016-0953-9.

21. Subramanian A, Tamayo P, Mootha VK, Mukherjee S, Ebert BL, Gillette MA, Paulovich A, Pomeroy SL, Golub TR, Lander ES, Mesirov JP. Gene set enrichment analysis: a knowledge-based approach for interpreting genome-wide expression profiles. *Proceedings of the National Academy of Sciences USA*. 2005;102(43):15545–15550. doi:10.1073/pnas.0506580102.

22. Liberzon A, Birger C, Thorvaldsdóttir H, Ghandi M, Mesirov JP, Tamayo P. The Molecular Signatures Database (MSigDB) hallmark gene set collection. *Cell Systems*. 2015;1(6):417–425. doi:10.1016/j.cels.2015.12.004.

23. Maslov S, Sneppen K. Specificity and stability in topology of protein networks. *Science*. 2002;296(5569):910–913. doi:10.1126/science.1065103.

24. Niwa H, Toyooka Y, Shimosato D, Strumpf D, Takahashi K, Yagi R, Rossant J. Interaction between Oct3/4 and Cdx2 determines trophectoderm differentiation. *Cell*. 2005;123(5):917–929. doi:10.1016/j.cell.2005.08.040.

25. Chazaud C, Yamanaka Y, Pawson T, Rossant J. Early lineage segregation between epiblast and primitive endoderm in mouse blastocysts through the Grb2-MAPK pathway. *Developmental Cell*. 2006;10(5):615–624. doi:10.1016/j.devcel.2006.02.020.

26. Rhinn M, Dollé P. Retinoic acid signalling during development. *Development*. 2012;139(5):843–858. doi:10.1242/dev.065938.

27. Tontonoz P, Hu E, Spiegelman BM. Stimulation of adipogenesis in fibroblasts by PPAR gamma 2, a lipid-activated transcription factor. *Cell*. 1994;79(7):1147–1156. doi:10.1016/0092-8674(94)90006-x.

28. Thiery JP, Sleeman JP. Complex networks orchestrate epithelial-mesenchymal transitions. *Nature Reviews Molecular Cell Biology*. 2006;7(2):131–142. doi:10.1038/nrm1835.

29. Pastushenko I, Brisebarre A, Sifrim A, Fioramonti M, Revenco T, Boumahdi S, Van Keymeulen A, Brown D, Moers V, Lemaire S, De Clercq S, Minguijón E, Balsat C, Sokolow Y, Dubois C, De Cock F, Scozzaro S, Sopena F, Lanas A, D'Haene N, Salmon I, Marine JC, Voet T, Sotiropoulou PA, Blanpain C. Identification of the tumour transition states occurring during EMT. *Nature*. 2018;556(7702):463–468. doi:10.1038/s41586-018-0040-3.

30. Bhatia S, Gunter JH, Burgess JT, Adams MN, O'Byrne K, Thompson EW, Duijf PHG. Stochastic epithelial-mesenchymal transitions diversify non-cancerous lung cell behaviours. *Translational Oncology*. 2023;37:101760. doi:10.1016/j.tranon.2023.101760.

31. Boyer LA, Lee TI, Cole MF, Johnstone SE, Levine SS, Zucker JP, Guenther MG, Kumar RM, Murray HL, Jenner RG, Gifford DK, Melton DA, Jaenisch R, Young RA. Core transcriptional regulatory circuitry in human embryonic stem cells. *Cell*. 2005;122(6):947–956. doi:10.1016/j.cell.2005.08.020.

32. Chakraborty M, Hu S, Visness E, Del Giudice M, De Martino A, Bosia C, Sharp PA, Garg S. MicroRNAs organize intrinsic variation into stem cell states. *Proceedings of the National Academy of Sciences of the United States of America*. 2020;117(12):6942–6950. doi:10.1073/pnas.1920695117.

33. Osorno R, Chambers I. Transcription factor heterogeneity and epiblast pluripotency. *Philosophical Transactions of the Royal Society B: Biological Sciences*. 2011;366(1575):2230–2237. doi:10.1098/rstb.2011.0043.

34. Chickarmane V, Enver T, Peterson C. Computational modeling of the hematopoietic erythroid-myeloid switch reveals insights into cooperativity, priming, and irreversibility. *PLoS Computational Biology*. 2009;5(1):e1000268. doi:10.1371/journal.pcbi.1000268.

35. Kageyama R, Ohtsuka T. The Notch-Hes pathway in mammalian neural development. *Cell Research*. 1999;9(3):179–188. doi:10.1038/sj.cr.7290016.

36. Jho EH, Zhang T, Domon C, Joo CK, Freund JN, Costantini F. Wnt/beta-catenin/Tcf signaling induces the transcription of Axin2, a negative regulator of the signaling pathway. *Molecular and Cellular Biology*. 2002;22(4):1172–1183. doi:10.1128/MCB.22.4.1172-1183.2002.

37. Katagiri T, Yamaguchi A, Komaki M, Abe E, Takahashi N, Ikeda T, Rosen V, Wozney JM, Fujisawa-Sehara A, Suda T. Bone morphogenetic protein-2 converts the differentiation pathway of C2C12 myoblasts into the osteoblast lineage. *The Journal of Cell Biology*. 1994;127(6 Pt 1):1755–1766. doi:10.1083/jcb.127.6.1755.

38. Kopan R, Ilagan MXG. The canonical Notch signaling pathway: unfolding the activation mechanism. *Cell*. 2009;137(2):216–233. doi:10.1016/j.cell.2009.03.045.

39. Kobayashi T, Mizuno H, Imayoshi I, Furusawa C, Shirahige K, Kageyama R. The cyclic gene *Hes1* contributes to diverse differentiation responses of embryonic stem cells. *Genes & Development*. 2009;23(16):1870–1875. doi:10.1101/gad.1823109.

40. Kawamura T, Suzuki J, Wang YV, Menendez S, Morera LB, Raya A, Wahl GM, Izpisúa Belmonte JC. Linking the p53 tumour suppressor pathway to somatic cell reprogramming. *Nature*. 2009;460(7259):1140–1144. doi:10.1038/nature08311.

41. Seale P, Sabourin LA, Girgis-Gabardo A, Mansouri A, Gruss P, Rudnicki MA. Pax7 is required for the specification of myogenic satellite cells. *Cell*. 2000;102(6):777–786. doi:10.1016/s0092-8674(00)00066-0.

42. Rocheteau P, Gayraud-Morel B, Siegl-Cachedenier I, Blasco MA, Tajbakhsh S. A subpopulation of adult skeletal muscle stem cells retains all template DNA strands after cell division. *Cell*. 2012;148(1–2):112–125. doi:10.1016/j.cell.2011.11.049.

43. Kuang S, Kuroda K, Le Grand F, Rudnicki MA. Asymmetric self-renewal and commitment of satellite stem cells in muscle. *Cell*. 2007;129(5):999–1010. doi:10.1016/j.cell.2007.03.044.

44. De Micheli AJ, Spector JA, Elemento O, Cosgrove BD. A reference single-cell transcriptomic atlas of human skeletal muscle tissue reveals bifurcated muscle stem cell populations. *Skeletal Muscle*. 2020;10(1):19. doi:10.1186/s13395-020-00236-3.
